# Comparative Proteomics Across Tissues and Crop Agroecosystems Reveals Agricultural Stressor Responses in the Western Honey Bee

**DOI:** 10.64898/2026.06.03.729970

**Authors:** Huan Zhong, Peipei Zhong, Junseo Park, Aleksandra Kozlova-Ryabova, Renata Moravcova, Jason C. Rogalski, Aidan Jamieson, Lance Lansing, Wendy W. T. Fang, Kyung-Mee Moon, Xiaojing Yuan, Lynae P. Ovinge, Jeff D. Kearns, Amanda S. Gregoris, Heather Higo, Julia Common, Ida M. Conflitti, Mateus Pepinelli, Lan Tran, Morgan Cunningham, Hosna Jabbari, Syed Abbas Bukhari, Sarah K. French, Jonathan Ho, Thomas B. Deckers, Jackie Zorz, Rodrigo Ortega Polo, Shelley E. Hoover, Stephen F. Pernal, Pierre Giovenazzo, Robert W. Currie, M. Marta Guarna, Amro Zayed, Leonard J. Foster

## Abstract

Maintaining honey bee health in crop production systems is increasingly difficult because worker bees encounter multiple chemical and biological pressures from pesticides and pathogens. How these field-realistic pressures affect molecular physiology across functionally distinct tissues remains poorly understood. Here, we tested whether tissue-resolved proteomics could separate stable tissue-specific patterns from crop-associated molecular changes. To do this, we profiled abdomen, gut, and head proteomes from honey bees collected across four Canadian crop ecosystems over two consecutive years, and integrated these data with pesticide-residue and pathogen-load measurements.

Proteomic variation was structured by both tissue identity and crop environment. Tissue-specific proteomic profiles were characterized across samples, whereas crop-associated effects were detected in both years and were stronger in 2021, the second year of the study. Tissue-specific enrichment and network analyses linked the abdomen to lipid catabolism and ubiquitin-proteasome proteostasis, the gut to central carbon metabolism, membrane transport, vesicle trafficking, and cytoskeletal organization, and the head to neurosensory and mitochondrial functions, together with amino-sugar metabolism and vesicle-associated quality-control modules. Among the measured pesticide residues, boscalid was the most reproducible chemical correlate of proteomic variation, with the strongest signal in the gut. Cross-year validation associated boscalid exposure with reduced abundance of gut proteins involved in mitochondrial metabolism, protein quality control, vesicle trafficking, nutrient transport, and biosynthetic pathways. Additionally, integrated proteome-transcriptome-microbiome factor analysis further identified gut-centered components associated with measured stressor variables and linked protein-level variation to coordinated transcriptomic and microbial shifts. Independent-year validation showed that compact crop-associated protein signatures detected in 2020 were also present in 2021. Together, these results show that honey bee tissues maintain stable proteomic identities while showing tissue- and year-specific responses to pesticide and pathogen pressures encountered in crop ecosystems. The gut proteome may specifically provide a sensitive molecular indicator of pesticide-associated perturbation under field conditions.

## Introduction

Western honey bees (*Apis mellifera*) are the most economically important managed pollinators in agricultural systems^1^. More broadly, animal-mediated pollination benefits approximately one-third of global crop production to some degrees^2^, reflecting the combined contribution of managed and wild pollinators. Widespread declines in colony health and increasing overwinter mortality rates have prompted efforts to identify the physiological mechanisms underlying colony loss^3–9^. Central to this effort is understanding how honey bee tissues maintain homeostasis under the diverse environmental pressures encountered during crop pollination.

Although every tissue of a honey bee shares an identical genome, individual tissues perform markedly distinct physiological roles that arise from differential gene expression and the assembly of specialised protein networks. For example, the abdominal fat body serves as a central metabolic and immune organ, integrating energy storage, detoxification, and humoral defence^10^. The midgut (hereafter "gut") mediates digestion and nutrient absorption while acting as a primary interface with ingested xenobiotics and pathogens.^11^ The head, which houses the brain and associated sensory structures, regulates neural signalling, behaviour, and environmental perception^12^. However, all current studies only focus on single tissues and/or controlled laboratory conditions^13–15^.

In agricultural landscapes, honey bees are simultaneously exposed to multiple classes of stressors whose composition and intensity vary across crop ecosystem and time. Chemically, bees encounter complex mixtures of agrochemicals through nectar, pollen, and hive matrices. Although typically present at sublethal levels, chronic exposure has been associated with impairments in energetically demanding processes, including reduced flight performance and altered reproductive output^16,17^. Additionally, bees are affected by biological stressors, such as multiple RNA viruses that disrupt host cellular homeostasis. Experimental infections have demonstrated widespread perturbations of host gene expression, including disruption of transcriptional regulation and protein synthesis pathways in infected bees^18^. These chemical and biological stressors co-occur in the field, and their relative contributions vary across crops and time, generating complex and composite exposure profiles^19–21^.

These stressors act through diverse physiological routes; therefore, their effects are unlikely to be uniform across tissues. Instead, crop-specific exposure environments may induce tissue-dependent changes in metabolic, immune, detoxification, digestive, and neural pathways^17,22–24^. A multi-tissue proteomic approach is therefore well suited to capture how physiological functions vary in honey bees under real-world pollination conditions.

In this study, we profiled the proteomes of three honey bee tissues (abdomen, gut, and head) collected from apiaries placed near and far from four crop types over two consecutive years (2020-2021): commodity canola (CAC), seed canola (CAS), cranberry (CRA), and highbush blueberry (HBB; **Figure 1**). For this purpose, we (1) identified proteins with consistent high expression across all crops within specific tissues to define tissue-stable core proteomes, (2) characterised environmentally sensitive proteins by assessing variation in expression across crops and quantifying dominant environmental influences using a meta-analytic approach and (3) isolated intrinsic protein-stressor relationships by applying constrained ordination and cross-year validated correlation analyses after accounting for crop-associated variation. Finally, we performed multi-omics analysis integrating proteomic, transcriptomic, and microbiome data to evaluate whether proteomic programs can be independently validated and to identify cross-omic coordination. This integrative analysis allows us to clearly see robust associations between specific stressors (e.g., *Vairimorpha* and VDV) and tissue-level molecular responses (e.g., suppression of ribosomal and translational machinery) in honey bee.

**Figure 1.**
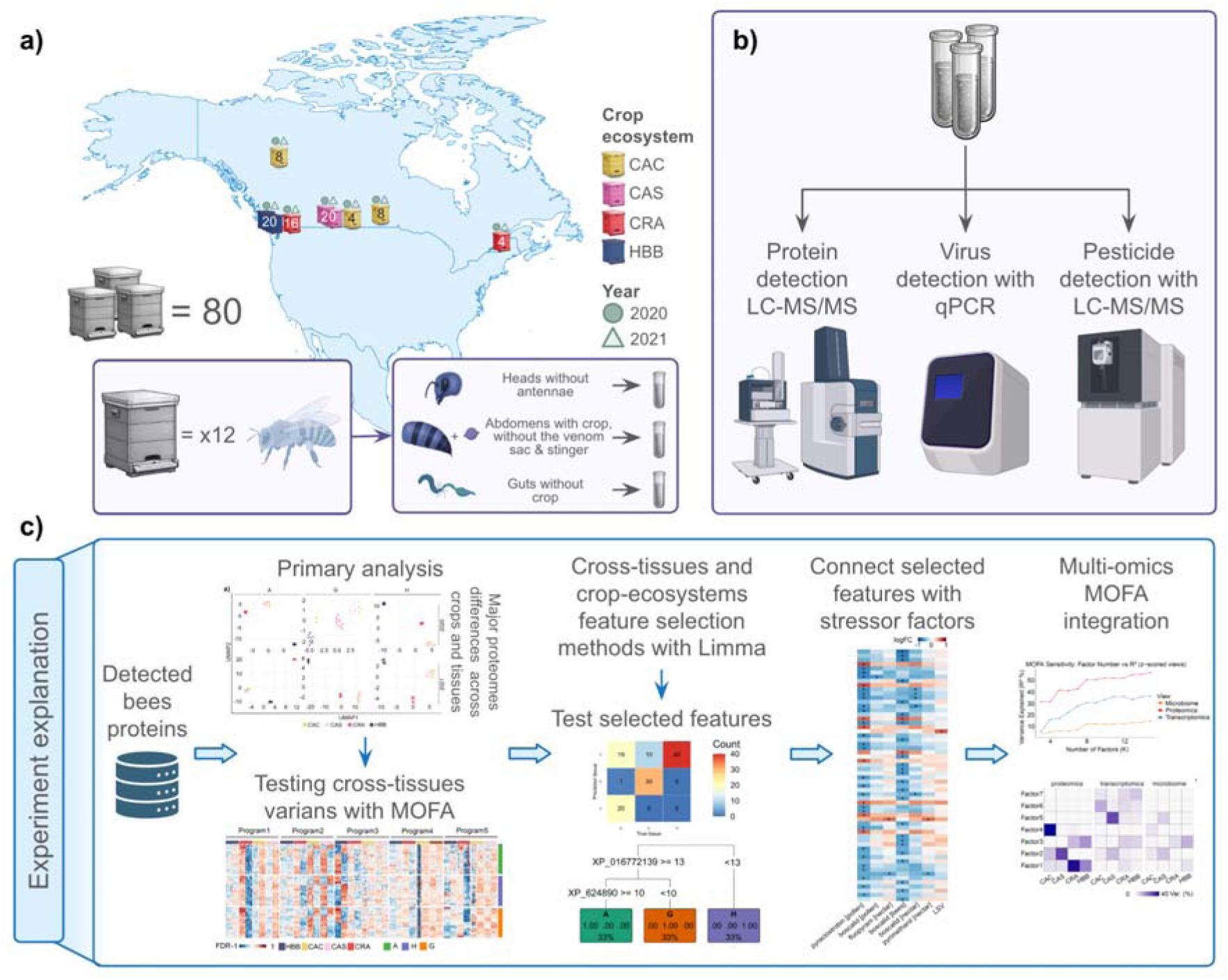
Overview of the experimental design and analytical workflow. (**a**) Sampling locations and colony collection. Each biological replicate represented one apiary comprising four colonies, from which nurse-aged worker bees were collected and dissected into head (H), abdomen (A; gut and stinger removed), and gut (G; honey crop removed, and for transcriptomics and proteomics only the midgut was processed) tissues. The numbers shown on the map correspond to the total number of colonies sampled per apiary, location, and crop type (CAC = commodity canola; CAS = seed canola; CRA = cranberry; HBB = highbush blueberry). In total, 80 colonies were studied across the 2020 and 2021 field seasons. Each apiary contained four colonies, from which three nurse bees per colony were collected. Bees were dissected into three tissue categories for proteomics analysis: the head (antennae removed), the abdomen (gut and stinger removed), and the gut. Dissections from 12 individuals were pooled within each apiary replicate and tissue type to generate a single sample, resulting in a total of 240 pooled samples ran in randomised order for proteomics analysis, as described in Zhong et al.^20^ **(b)** Molecular measurements including proteomics (LC-MS/MS), virus quantification (qPCR), and pesticide detection (LC-MS/MS). **(c)** Proteomic and computational analysis workflow, including differential abundance analyses, feature selection, stressor association analyses, and multi-omics integration using MOFA.

## Materials and Methods

### Sample collection and crop-system grouping

To conduct the experiment, adult nurse-aged worker bees were collected from apiaries associated with four crop-system contexts: Commodity canola (CAC), Seed canola (CAS), Cranberry (CRA), and Highbush blueberry (HBB). CAC represented commodity canola production within prairie crop rotations in Western Canada, whereas CAS represented hybrid seed canola production, a more specialized system concentrated in irrigated areas of southern Alberta. CRA and HBB represented perennial fruit-crop systems with distinct pesticide-exposure profiles and were sampled across sites in British Columbia and Québec (**Figure. 1a**).

In each of the 2020 and 2021 field seasons, apiaries were sampled across 40 sites, with ten apiary-level replicates per crop system; some sites were sampled in both years (**Figure. 1a**). Within each crop system, apiaries included sites located near the focal crop and paired sites located farther from it. Because previous work in British Columbia showed that honey bees can encounter pesticide residues at both near and far sites relative to highbush blueberry fields^25–27^, near and far apiaries were pooled within each crop group for the primary analyses. To evaluate whether this pooling affected the interpretation of global proteomic structure, we performed a supplementary near/far sensitivity analysis within each crop-by-tissue subset after year-wise adjustment. Near/far separation was quantified using silhouette scores in the global protein expression space and visualized using PCA ordinations. Crop group was therefore treated as a crop-associated agroecosystem variable, rather than as the isolated effect of a single crop species, because crop identity was partially aligned with geographic region and beekeeper management practices.

Each apiary replicate comprised four colonies. From each colony, three nurse-aged worker bees were collected and dissected into head, abdomen, and gut tissues. The head was processed after antennae removal, the abdomen after removal of the gut and stinger, and the gut after removal of the honey crop. Tissue dissections from 12 bees per apiary were pooled to generate one proteomic sample per tissue and apiary. This design yielded 120 proteomic samples per year across four crop-system contexts, three tissues, and ten apiary-level biological replicates per crop system.

### Proteomics Processing

Proteins from three tissue types were extracted by mechanical homogenization in a lysis buffer containing 4% sodium dodecyl sulfate (SDS) and 100 mM tris(hydroxymethyl)aminomethane hydrochloride (Tris-HCl), pH 6.8, supplemented with protease inhibitors. Protein concentration was measured using a bicinchoninic acid (BCA) assay. For downstream processing, 40 µg of protein from head and abdomen samples and 60 µg from gut samples were reduced with dithiothreitol (DTT), alkylated with chloroacetamide (CAA), purified by sodium dodecyl sulfate-polyacrylamide gel electrophoresis (SDS-PAGE), and subjected to in-gel 0.45 µg tryptic digestion. Resulting peptides were desalted using homemade C18 STAGE-tips^28^, quantified by absorbance at 205 nm, and 150 ng per sample was analyzed on a NanoElute UHPLC system coupled to a timsTOF Pro 2 mass spectrometer operated in DIA-PASEF mode. Raw data were searched using DIA-NN within FragPipe^29^ against a previously generated spectral library, with additional acquired sites searched alongside the main dataset but excluded from downstream analysis. A more comprehensive description of the proteome extraction and LC-MS/MS analysis procedures is provided in the Supplementary Methods^20^.

### Sample collection for pathogen and agrochemical measurements

For pathogen and agrochemical-residue measurements, a separate batch of adult worker bees was sampled from honey frames in the same colonies used for proteomic sampling. From each colony, 15 workers were pooled for pathogen quantification and agrochemical-residue analysis, following Zhong et al.^20^ (**Supplementary Figure. 1**). These samples were collected during the crop-specific mid-bloom sampling period and provided matched colony- or apiary-level stressor profiles for integration with the tissue proteomic data.

### Agrochemical-residue analysis

Agrochemical residues were measured by LC-MS/MS across up to four matrices: adult bees, nectar, pollen and wax. The screening panel covered 50 unique pesticide compounds, corresponding to 132 compound-matrix combinations. Six pesticide compounds were detected in more than 30% of samples. The complete compound list, sample matrix, chemical class, units, detection frequency and compound- or matrix-specific detection or quantification limits are provided in **Supplementary Table 1**. For downstream stressor-association analyses, pesticide variables were retained only when they showed sufficient detection frequency and sample coverage across crop-year groups.

### Viral and Vairimorpha analysis

Viral infection was assessed from pooled adult worker-bee samples by RT-qPCR, following Alison et al.^21^ Nine honey bee viruses were quantified: *Deformed wing virus* (DWV), *Chronic bee paralysis virus* (CBPV), *Acute bee paralysis virus* (ABPV), *Kashmir bee virus* (KBV), *Varroa destructor virus* (VDV), *Sacbrood virus* (SBV), *Black queen cell virus* (BQCV), *Israeli Acute Paralysis Virus* (IAPV) and *Lake Sinai virus* (LSV). Viral titres were reported as copies per bee. Six viruses were detected in more than 30% of samples and were retained for downstream summary and association analyses. *Vairimorpha spp.* burden was measured from the same pooled adult worker-bee samples, following Alison et al.^21^ *Vairimorpha* load was reported as spore load per bee and retained as a pathogen-burden variable when sample coverage was sufficient.

For visualization and modelling, pesticide-residue, viral-load and *Vairimorpha*-load values were log_10_ (x+1) to accommodate non-detects and reduce right-skewed distributions. For composite burden variables, x represents the summed load within the relevant compound, matrix or pathogen category. Group-level median loads were then computed for each crop-year group. Virus prevalence was defined as the proportion of samples within each crop-year group with detectable viral copy number (>0). Median-load and prevalence summaries were visualized as heatmaps.

### Data preprocessing and preliminary analysis

All analyses were carried out in R (v4.4.2). Raw protein abundance data were obtained by label-free quantitative (LFQ) mass spectrometry. Contaminant proteins were removed based on a custom UniProt contaminant list, predefined prefixes (e.g., CON_), and keyword filtering (e.g., keratin, trypsin). Phenotypic data were restricted to samples collected at mid-bloom (timepoint 2) in both sampling years. This yielded 120 samples per year representing four crop environments (CAC, CAS, CRA, and HBB) and three tissues: abdomen (A), gut (G), and head (H). 12 experimental groups per year with ten biological replicates per group (Crop×Tissue×Year) were generated. All subsequent preprocessing steps were performed independently for each sampling year to preclude cross-year data leakage.

Protein-wise missingness was calculated within each experimental group, and proteins were processed using a hybrid strategy: proteins with no missing values were retained, proteins with ≤60% missingness were imputed using missForest^30^, and proteins with >60% missingness were excluded from the corresponding group. Protein abundances were logL(x + 1)-transformed before downstream analyses.

After imputation, multivariate proteomic structure was assessed using PCA and UMAP^31^ on centered and scaled protein abundance matrices, with zero-variance proteins removed. Group-level differences were tested using PERMANOVA^32^ on Bray-Curtis dissimilarities, with homogeneity of dispersion assessed using betadisper. Differential protein abundance was analyzed with limma using empirical Bayes moderation^33^, followed by Benjamini-Hochberg FDR correction^34^. Proteins with |logL fold change| ≥ 1 and FDR < 0.05 were considered significantly differentially expressed. Functional enrichment of up- and down-regulated proteins was performed using clusterProfiler for GO terms and KEGG pathways^35–37^. A detailed description of missing-value handling, multivariate analyses, differential expression testing, and enrichment analysis is provided in the Supplementary Methods.

### Multi-layer Categorisation and Filtration Framework

A multi-layer categorization framework was applied to identify proteins with tissue-specific and crop-responsive expression patterns. First, tissue specificity was assessed using pairwise tissue contrasts, requiring consistent significant changes in contrasts involving the target tissue and minimal variation in the background contrast. Proteins were classified as tissue-specific, tissue-conserved, or tissue-partial, with cross-year validation requiring the same label in both 2020 and 2021. Tissue-specific protein sets from both years were further analyzed using STRING^35^ to assess protein-protein interaction enrichment, functional annotation, network clustering, and hub protein identification. To characterize crop-related proteomic responses, differential expression results were integrated across three increasingly stringent layers: block-level classification of crop-contrast signals, protein-level identification of tissue-specific crop sensitivity, and random-effects meta-analysis^36^ to estimate crop-dominant effects within each tissue and year. Pooled crop ecosystem effects were modeled using metafor, and dominant crops were defined by the largest significant absolute pooled effect.

### Analysis of Stressor-Proteome Interactions

Stressor-proteome interactions were assessed using distance-based redundancy analysis^37^ (db-RDA), in which tissue-specific protein abundance matrices were constrained by the stressor matrix. Crop-related batch effects were removed from the 2020 protein and stressor data, and the same 2020-derived correction parameters were applied to the 2021 dataset for validation. Protein matrices were centered and scaled, and zero-variance features were removed. Model significance was evaluated by permutation ANOVA, and proteins associated with the constrained ordination space were shortlisted using permutation-based vector fitting.

For each tissue, shortlisted proteins were tested for associations with stressors using Spearman correlations in the 2020 dataset, followed by FDR correction. Significant protein-stressor pairs were then evaluated in the independent 2021 dataset using a regression-based reproducibility test that assessed both direction consistency and statistical significance. To reduce crop-driven confounding, stressors with high crop-associated variance were excluded based on the intraclass correlation coefficient. Protein-stressor associations were considered reproducible if they were significant in 2020, validated in 2021, and not strongly confounded by crop identity.

### MOFA Analysis

Multi-omics factor analysis was performed using MOFA2^38^ to identify latent sources of variation across tissues and omics layers. Protein abundance data were logL-transformed, filtered for missingness and zero-variance features, and z-score standardized. For the tissue-view analysis, each tissue was treated as a separate view and crop as a group. For the integrated gut multi-omics analysis, only samples with matched proteomics, transcriptomics, and microbiome data were retained. RNA-seq counts were normalized to logL(CPM + 1), microbiome data were CLR-transformed, and all omics matrices were variance-filtered and standardized before integration.

Two MOFA2 models were trained: one to compare proteomic variation across tissues and another to integrate gut proteomics, transcriptomics, and microbiome profiles. Model dimensionality was optimized by evaluating the variance explained across different numbers of latent factors. Factor loadings and crop-wise factor scores were used to identify molecular features contributing to crop-specific responses. Associations between latent factors and environmental stressors were tested using Spearman correlations with FDR correction. Functional interpretation of factor-associated proteins and genes was performed using GO and KEGG enrichment analyses.

### Analysis design for temporal reproducibility of crop-associated molecular profiles

To evaluate whether crop-associated molecular profiles were reproducible across field seasons, we applied a temporal validation framework in which model development and independent testing were separated by year. Samples collected in 2020 formed the training cohort used for supervised feature selection, model optimisation, and classifier fitting. The resulting feature panels were then locked and applied without modification to samples collected in 2021, which served as an independent validation cohort.

Analyses were performed separately for abdomen, gut, and head tissues, with crop identity treated as a four-class outcome. Sparse molecular signatures were selected using an initial XGBoost-based screening step followed by recursive feature elimination with cross-validation. Protein-only, RNA-only, and combined RNA + protein models were evaluated separately. For the combined models, features were selected within each molecular layer before being concatenated into an early-fusion matrix. Final selected feature panels were evaluated using tuned Random Forest classifiers, and model performance in the independent 2021 cohort was summarised using one-vs-rest multiclass metrics, including AUC, precision, recall, and F1-score.

UMAP embeddings were used as descriptive summaries of sample structure within each tissue and feature set. To quantify whether molecular separation was more strongly associated with crop type than with mapped sampling location, we calculated a crop-location structure score:

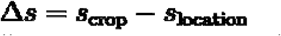

where s_crop and s_location are silhouette scores computed on the two-dimensional UMAP coordinates using crop and mapped-location labels, respectively^39^. Positive values indicate stronger crop-associated than location-associated separation.

### Evaluation metrics

Macro-average AUC was used to summarise class-balanced discrimination across crop classes^40^:

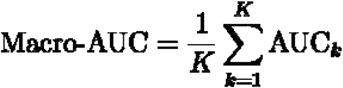

where K is the number of crop classes and AUC_k is the one-vs-rest AUC for crop class k. Micro-average AUC was calculated by pooling all one-vs-rest class decisions before computing the AUC^40^:

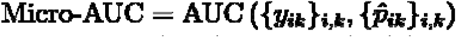

Precision, recall, and F1-score were reported as class-averaged validation metrics for the independent 2021 cohort.

## Results and Discussion

To establish the global architecture of proteomic variation, we first assessed data quality, ordination patterns, and the relative contributions of tissue and crop to overall variance in samples collected in the different crop agroecosystems (**Figure. 1**). Label-free quantitative mass spectrometry of 240 samples (120 per year; 4 crops × 3 tissues × 10 biological replicates) yielded a total of 5815 and 5852 quantified proteins in 2020 and 2021, respectively, after quality filtering. Mean per-sample detection counts were consistent between years and across the four crops (commodity canola [CAC], seed canola [CAS], cranberry [CRA], and highbush blueberry [HBB]), with all crop-level means falling within a 10% range (**Table 1, Supplementary Figure. 1a**). Standard deviations remained low across all groups (**Table 1**). Detection missingness was also comparable between years, with 23.1% (2020) and 22.2% (2021) of protein × sample observations recorded as missing. At the protein × group level, missingness showed a polarised distribution^41^: the majority of entries (61-64%) had no missing values, whereas a smaller subset of presumably low-abundance proteins (13.5-14.0%) displayed >80% missingness within at least one experimental group. After imputation and complete-case filtering, which required no missing values per protein and nonzero variance, 3722 proteins remained in 2020 and 3689 in 2021, with 3470 proteins shared between years. Because near and far apiaries were pooled within crop groups for the primary analyses, we next performed a sensitivity analysis to determine whether apiary placement represented a dominant axis of proteomic variation. Near/far separation was assessed in the global protein expression space within each crop-by-tissue subset after year-wise adjustment. Silhouette scores remained consistently low across crop-by-tissue subsets, and PCA ordinations showed substantial overlap between near and far samples. These results indicate that near/far placement contributed only weakly to global proteomic variation and did not define a dominant separable subgroup **(Supplementary Figure 2)**.

**Table 1.**
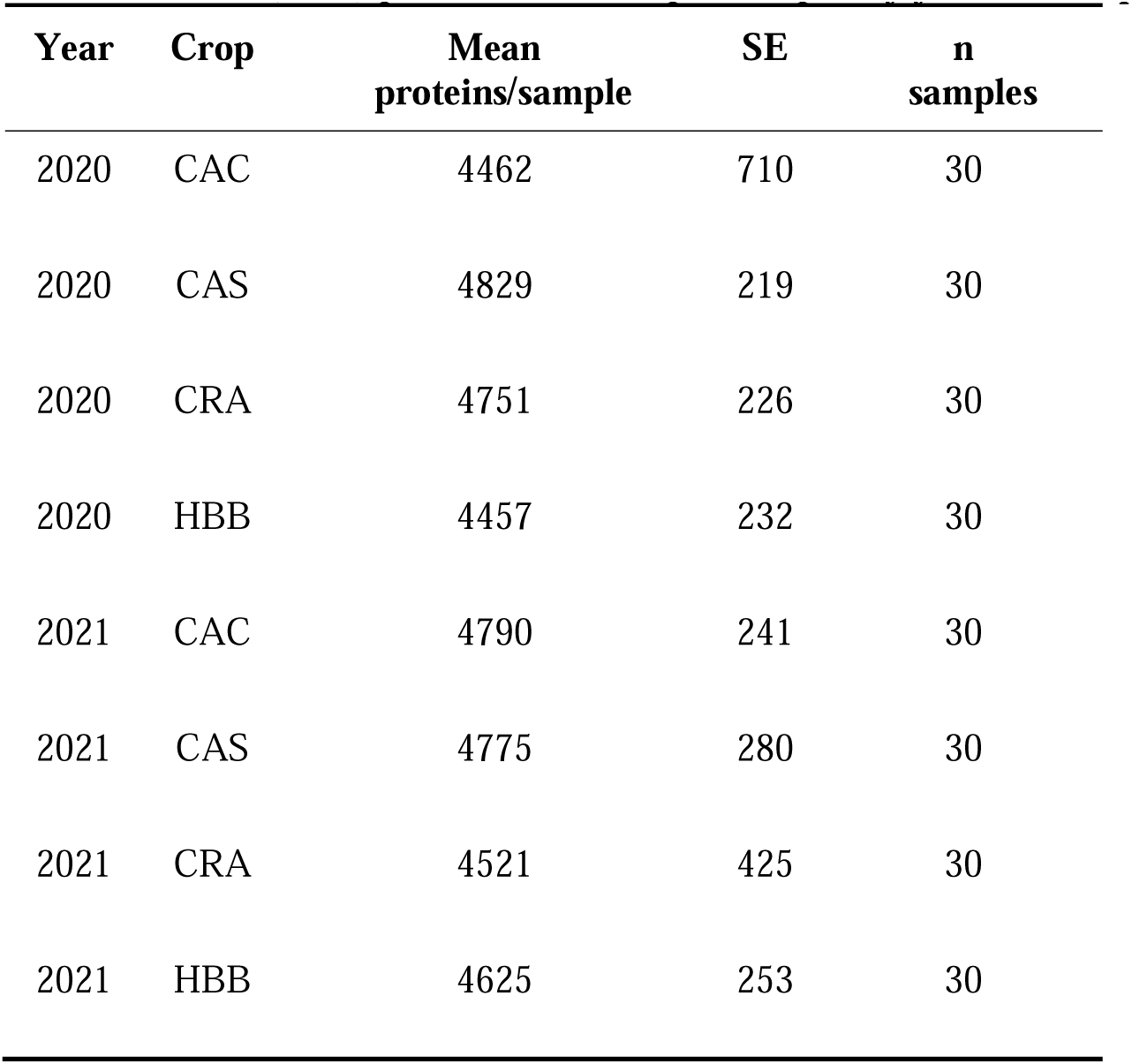
Mean (± SD) proteins detected per sample, by year and crop.

| Year | Crop | Mean<br>proteins/sample | SE | n<br>samples |
| --- | --- | --- | --- | --- |
| 2020 | CAC | 4462 | 710 | 30 |
| 2020 | CAS | 4829 | 219 | 30 |
| 2020 | CRA | 4751 | 226 | 30 |
| 2020 | HBB | 4457 | 232 | 30 |
| 2021 | CAC | 4790 | 241 | 30 |
| 2021 | CAS | 4775 | 280 | 30 |
| 2021 | CRA | 4521 | 425 | 30 |
| 2021 | HBB | 4625 | 253 | 30 |

### Global cross-tissue MOFA resolves crop- and stressor-linked axes of proteomic variation

Global proteomic variation in field-exposed bees arises from the intersection of tissue specialization and crop environment. Head, gut and abdomen provide distinct physiological contexts, while each crop system carries a different combination of pesticide residues and pathogen burden. We therefore used MOFA2 to analyse the three tissues together, resolving proteome-wide variation into latent programs that captured tissue contribution, crop activity and cross-tissue covariance linked to measured stressors (**Figure. 2a**). Each latent factor (hereafter referred to as a program for single proteomics by MOFA) was characterised by (i) variance explained across tissues (**Figure. 2b-c**), (ii) protein loading weights defining protein contributions, and (iii) program scores that can be correlated with measured stressors.

**Figure 2.**
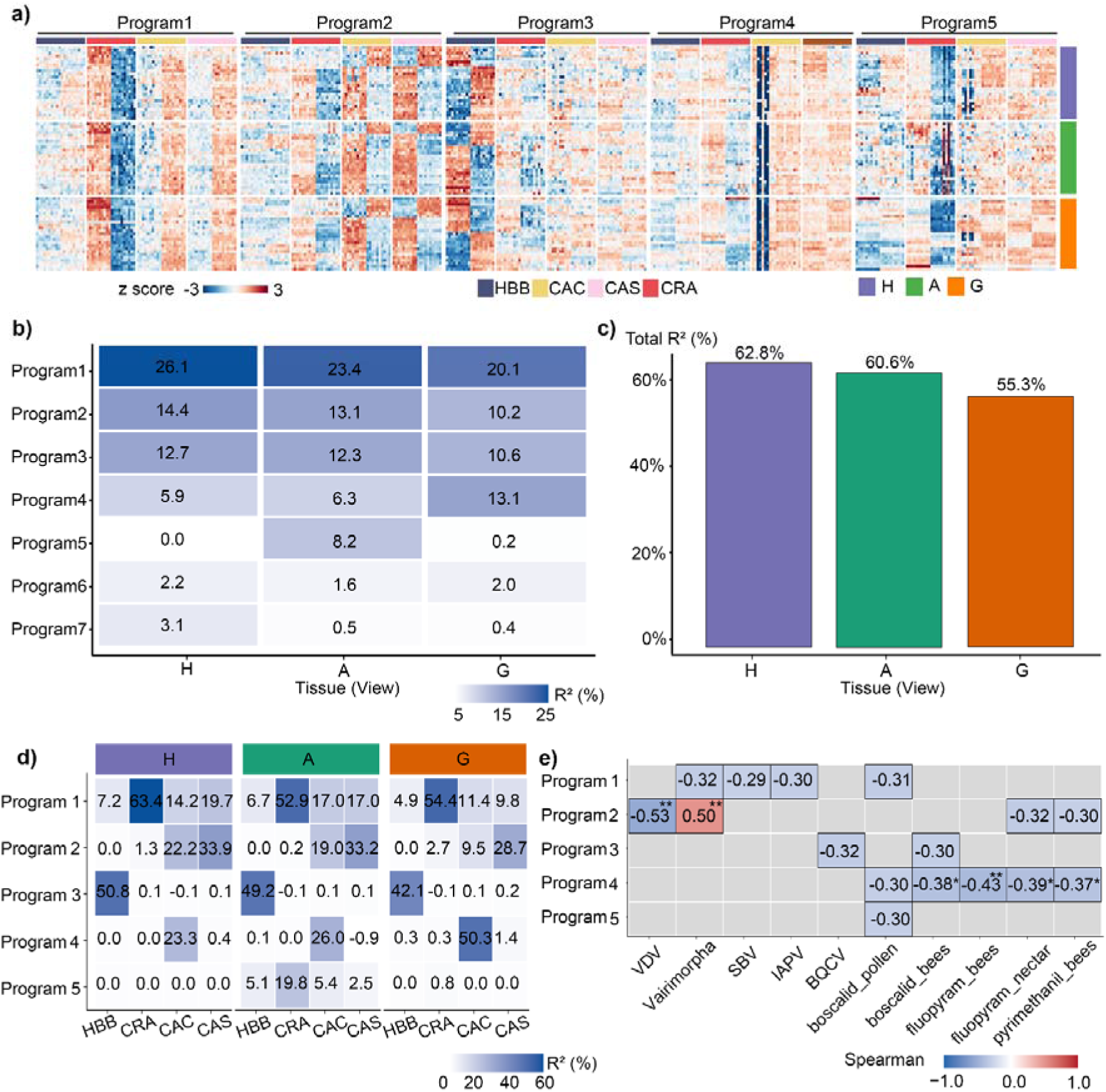
Cross-tissue covariance patterns with different activities in each crop for the proteome dataset based on MOFA. (**a**) Z-scored protein abundance across samples within each tissue. Colors represent standardized protein abundance (z-scores) computed across samples for each protein within each tissue (head [H], gut [G], abdomen [A]). Red indicates higher-than-average abundance and blue indicates lower-than-average abundance relative to the protein-specific mean. Each row represents a protein grouped by tissue of origin. Columns represent samples ordered by crop groups (CAC = commodity canola; CAS = seed canola; CRA = cranberry; HBB = highbush blueberry). **(b)** Variance explained (R^2^) by each MOFA latent program across tissues, averaged across crops. **(c)** Total variance explained by all latent programs within each tissue. **(d)** Variance explained by each latent program across crops within each tissue, showing tissue-specific program activity patterns. **(e)** Associations between latent programs and biotic or chemical stressors. Positive and negative associations are shown in red and blue, respectively. Significance levels are indicated by *FDR < 0.05 and **FDR < 0.01, with non-significant associations shown in grey. The sign of each association indicates whether stressor exposure is associated with higher (positive) or lower (negative) program scores.

Program 1 explained the largest share of total variance across all four crops. Its effect was overwhelmingly concentrated in CRA, which captured 52.9-63.4% of variance across the three tissues. However, variance explained in HBB, CAC, and CAS (**Figure. 2d**) was substantially lower. Program 1 scores were negatively correlated with boscalid in pollen (ρ = -0.314), *Vairimorpha* spore load (ρ = -0.316), IAPV viral titre (ρ = -0.305), and SBV (**Figure. 2e**). Proteins (with negative loadings on Program 1) were enriched for chromatin- and translation-associated functions in the gut. The top enriched terms were "structural constituent of chromatin" and "nucleosome", followed by "signaling receptor activity", "protein heterodimerization activity", and "large ribosomal subunit". A parallel chromatin enrichment was observed in head tissue (**Supplementary Figure. 3**). Program 2 captured variance predominantly in CAS and CAC, reflecting the shared stressor association with VDV (ρ = - 0.535), followed by *Vairimorpha* (ρ = +0.503), fluopyram in nectar and pyrimethanil in bees. Notably, VDV and *Vairimorpha* showed opposing correlation directions (**Figure. 2e**). Enrichment of the positively weighted proteins revealed oxidative phosphorylation in abdomen and translation-related terms in gut, including "structural constituent of ribosome" and "mitochondrial outer membrane". Proteins with negative loadings were enriched for galactose metabolism and nucleotide sugar biosynthesis in the gut (**Supplementary Figure. 3**). Program 3 was sharply restricted to HBB, explaining 42.1-50.8% of variance across all tissues. The associated stressors were BQCV (ρ = -0.319) and boscalid in adult bees (ρ = -0.304; **Figure. 2e**). Enrichment analysis of the negatively loaded proteins identified integrin signalling in head and a cytoplasmic localisation term in abdomen as the top functional signatures (**Supplementary Figure. 3**). Program 4 was concentrated in CAC (23.3-50.3% across tissues, with gut showing the highest R^2^). It showed the strongest and most numerous associations with pesticide exposure among all programs: fluopyram in bees and nectar, boscalid in bees and pollen, and pyrimethanil in bees (ρ = -0.301 to -0.430; **Figure. 2e**). The enrichment profile of negatively loaded proteins was dominated by translational machinery in head tissue: "ribosome", "translation", "structural constituent of ribosome", and the KEGG ribosome pathway. In the gut, proteolysis-related terms were also enriched among negatively loaded proteins. In contrast, positively loaded proteins in the head were enriched for microtubule cytoskeleton organisation (**Supplementary Figure. 3**). Program 5 was the only tissue-restricted program, driven exclusively by abdomen across all crops, with CRA again showing the highest contribution (CRA = 19.8%, HBB = 5.1%, CAC = 5.4%, CAS = 2.5%). The sole significant stressor association was boscalid in pollen (ρ = -0.299; **Figure. 2e**). Positively loaded proteins were enriched for metalloendopeptidase activity and cullin-RING ubiquitin ligase complex in abdomen (**Supplementary Figure. 3**). Together, these results show that proteome-wide variation was organized into separable global axes rather than a single uniform crop or tissue effect.

**Figure 3.**
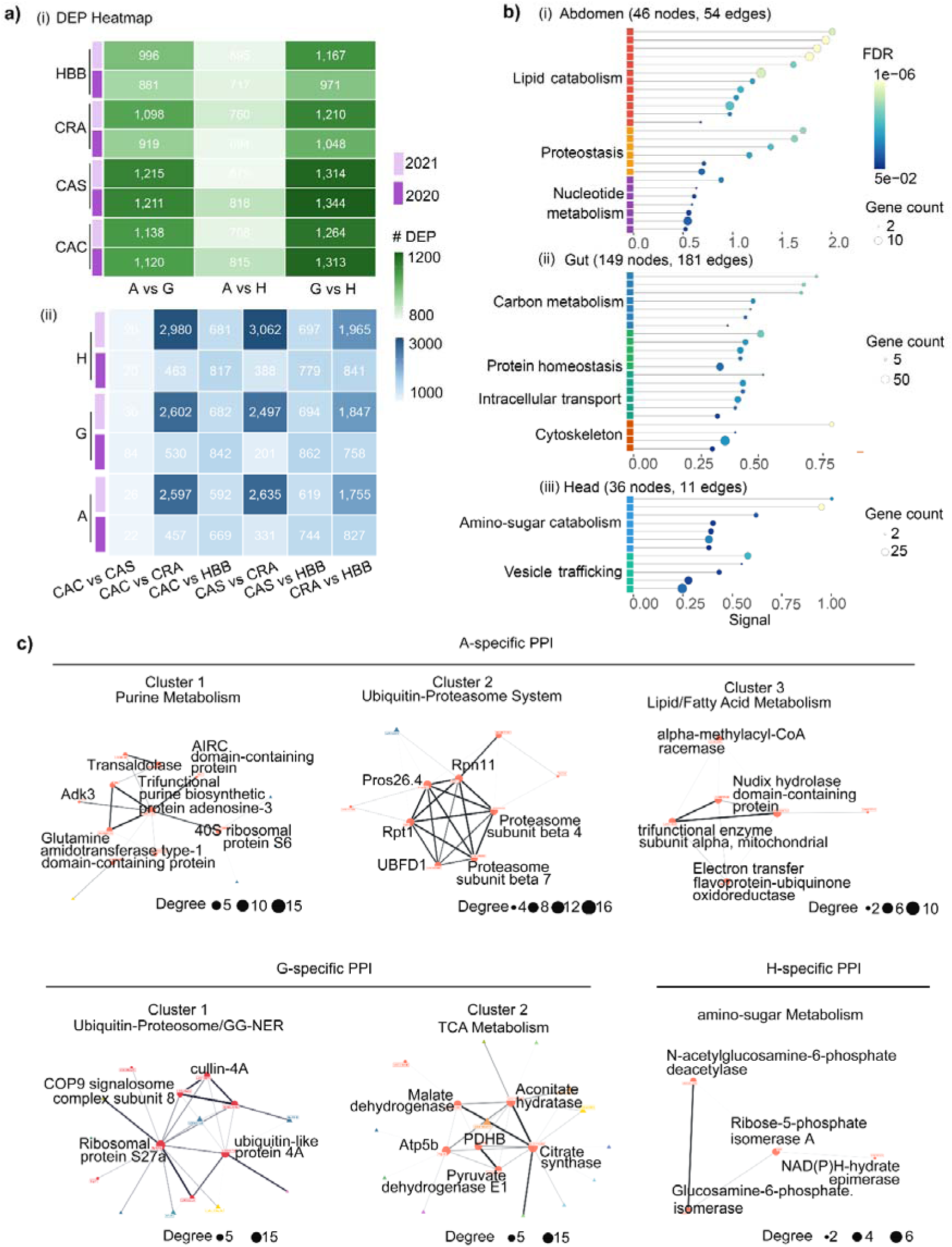
Tissue-stable core proteomes of honey bee head (H), abdomen (A), and gut (G). (**a**) Numbers of differentially expressed proteins (DEPs) identified in tissue(i) and crop (ii) contrasts across 2020 and 2021. CAC = commodity canola; CAS = seed canola; CRA = cranberry; HBB = highbush blueberry. **(b)** Functional enrichment analysis of tissue-specific proteins in abdomen (i), gut (ii), and head (iii) tissues. Bubble size represents gene count, and colour indicates false discovery rate (FDR). Complete GO enrichment results are provided in Supplementary Table 2, 3, 4; Supplementary Figure 2c. **(c)** Representative protein-protein interaction (PPI) subnetworks derived from tissue-specific proteins. Nodes represent proteins and edges indicate known or predicted interactions. Highlighted clusters correspond to enriched functional modules within each tissue. Networks were generated using the STRING database (v12.0) with a confidence score cutoff of >0.4.

### Proteomic variation was structured primarily by tissue identity, with crop-associated effects differed in two years

We then moved from global programs to individual proteins, in order to determine whether the latent tissue and crop programs were reflected in individual protein changes. Protein expression matrices varied significantly across crop and tissue in both years (PERMANOVA, p = 0.001 for all terms; **Table 2**). Crop explained 22.6% of variance in 2020 and 49.8% in 2021, whereas tissue explained 47.7% and 35.1%, respectively. This indicated a relative strengthening of cropLassociated variation. Dispersion in protein expression levels differed across the four crops (**Table 2, Supplementary Figure. 1b**). In 2020, CAC showed the broadest spread (median distance to centroid = 0.019) and CRA in 2021 (0.020), while HBB remained the most compact in both years.

**Table 2.**
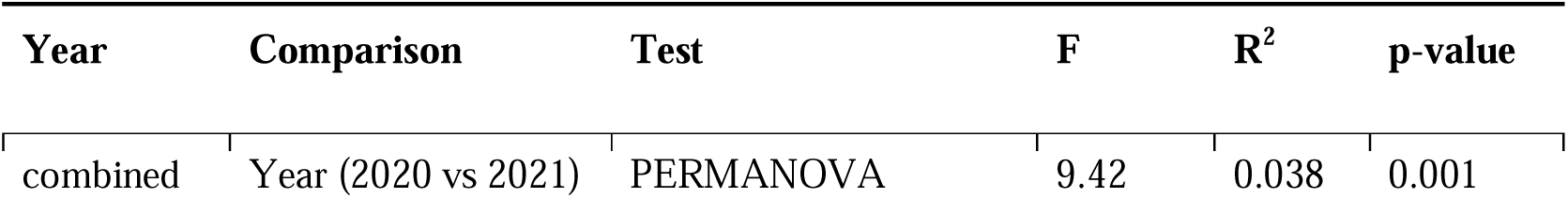
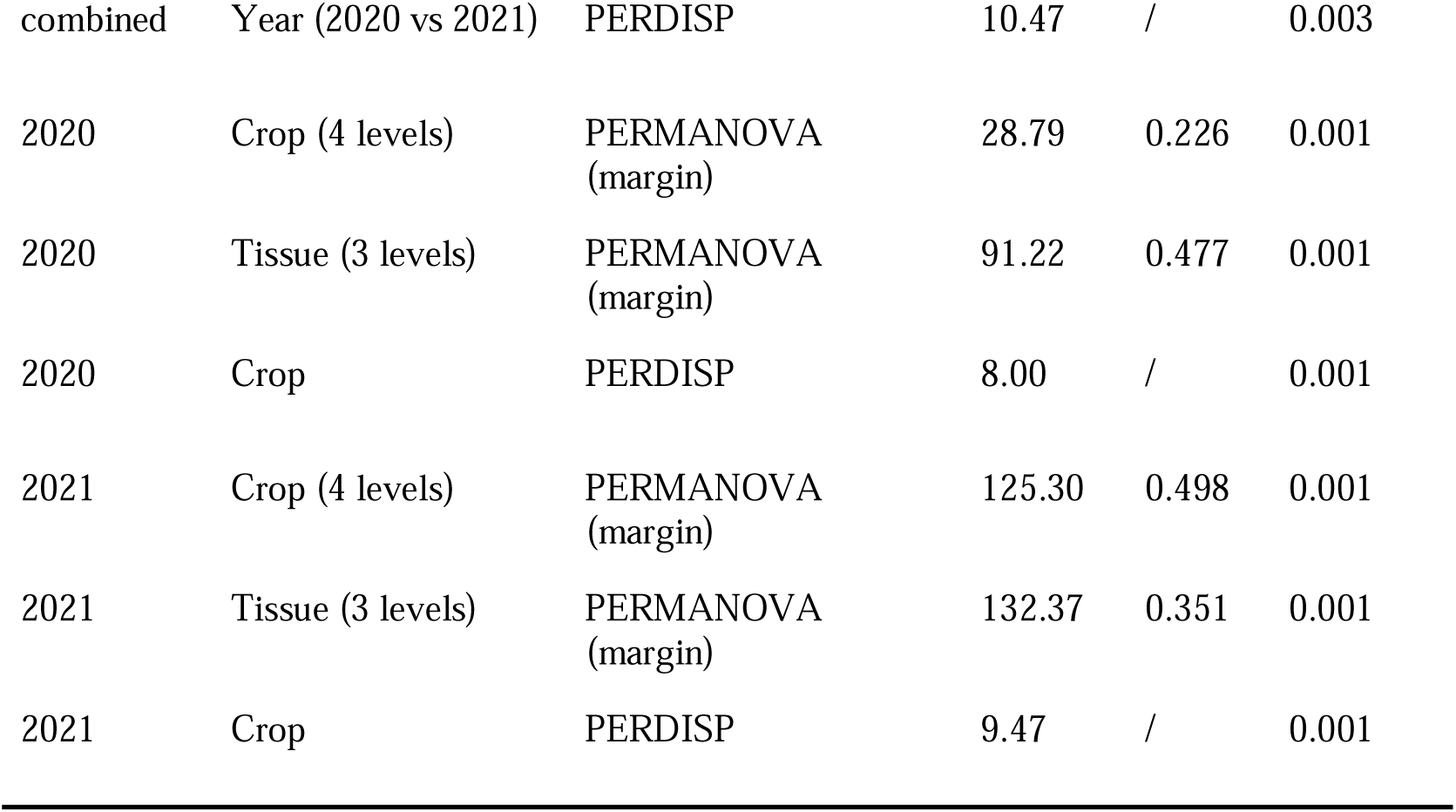
PERMANOVA and PERDISP tests of crop and year effects on protein composition (Bray-Curtis dissimilarity, 999 permutations).

To visualize these patterns, we performed UMAP, computed separately for each tissue to avoid tissue-driven variation. CAC and CAS consistently overlapped, whereas CRA and HBB formed distinct clusters (**Supplementary Figure. 1c**). Principal component analysis (PCA) on centered and scaled expression data visually supported this trend (**Supplementary Figure. 1d**). In 2020, the first two principal components (PCs) captured 21.6% (PC1) and 17.8% (PC2) of total variance in protein expression. In 2021, PC1 increased to 40.6% and crops showed clearer separation, particularly CRA. It formed a distinct cluster along PC1, which was consistent with the higher crop ecosystem effect observed in PERMANOVA.

Differential expression analysis across 60 pairwise comparisons revealed similar hierarchical and contextLdependent organisation of proteomic variation. Tissue contrasts consistently yielded 679-1344 differentially expressed proteins across two years (DEPs; **Figure. 3a**). Within tissues, a conserved pattern of relative divergence was observed in all crops and years: the gut-head contrast yielded the largest number of DEPs, followed by abdomen-gut and abdomen-head comparisons. Crop contrasts showed a wider range of DEP counts (**Figure. 3a**). Pairwise comparisons among nonLCRA crops (e.g., CAC vs CAS) yielded 20-150 DEPs in both years for each tissue. In contrast, all comparisons involving CRA produced ∼300 DEPs in 2020 (e.g., CAS vs CRA), whereas in 2021, these same contrasts generated ∼3000 DEPs for each tissue. Together, tissue identity defined a stable baseline of proteomic variation across years and crops. However, crop ecosystem effects can induce substantial shifts that, in some cases, equal or exceed the magnitude of tissue-driven divergence, yet the underlying tissue organisation remains detectable as a consistent secondary axis.

### Tissue identity defines stable core proteomes with distinct functional programs

Having established that tissue identity is the dominant and most stable axis of proteomic variation (accounting for 35-48% of variance across years), we next sought to characterize the functional programs that underpin each tissue’s molecular identity. Each tissue exhibited a distinct, coherent functional program across both years (**Supplementary Figure. 4a**). Head tissue was consistently enriched for neurosensory signaling, mitochondrial energy metabolism, and active protein synthesis, as evidenced by KEGG pathways including “Phototransduction - fly”, “Neuroactive ligandLreceptor interaction”, and “Oxidative phosphorylation”, as well as GO terms “signal transduction” and “potassium ion transport”. Additional enrichment of mitochondrial electron transport chain, ATP synthesis, and ribosomeLrelated terms (e.g., structural constituent of ribosome, translation) supported the high energetic and translational demand of neural tissue.

Gut tissue was enriched for processes related to protonLdriven membrane transport, acidification, and oxidoreductive metabolism. Representative GO terms included “protonLtransporting ATPase activity” and “proton transmembrane transport”, as well as KEGG pathways “Lysosome” and “Phagosome”, indicating acidificationLdependent intracellular processing. Additional enrichment of “monooxygenase activity”, “heme binding”, “Pentose phosphate pathway”, and “Folate biosynthesis” suggested its potential roles in detoxification and redox metabolism (**Supplementary Figure. 4a**).

Across tissues, abdomen showed the highest mean -log_10_(FDR), with enriched terms primarily related to proteostasis and systemic metabolism, especially for a ubiquitinLproteasome module (GO: “proteasome core complex”, “threonineLtype endopeptidase activity”; KEGG: “Proteasome”) and pathways of lipid and amino acid catabolism (“Fatty acid metabolism”, “One carbon pool by folate”, and “Valine, leucine and isoleucine degradation”, **Supplementary Figure. 4a**).

These enrichment profiles were consistent with the characteristic physiological functions of each tissue. However, they were derived from all DEPs identified in pairwise tissue contrasts. Proteins involved may just be generally higher in one tissue, rather than the conserved features defining tissue identity. To further resolve the core molecular architecture underlying tissue-specific functional programs, we applied a more stringent categorization on DEPs to identify proteins that were significantly and consistently enriched within each tissue across all four crops and both years (**Supplementary Figure. 5**). The tissue-specific proteins in each year were input for PPI network analysis (**Supplementary Figure. 6**). Among abdomen-enriched proteins, 30 candidates were identified in 2020 and 27 in 2021, of which four were retained in both years (13.3% of the 2020 set). Gut showed stronger conservation, with 41 of 86 candidates retained across years despite a larger 2021 set of 170 proteins (47.7%). Head showed more limited conservation, with two of 16 candidates retained across years, compared with 33 candidates in 2021 (12.5%).

### Abdomen maintained metabolic homeostasis through coupled lipid catabolism and proteasome-mediated proteostasis

The abdomen-specific PPI network revealed the most tightly integrated architecture among the three tissues, as indicated by the lowest PPI enrichment p value and highest clustering coefficient (PPI enrichment p = 7.44×10^-15^, clustering coefficient = 0.572; **Table 3**). MCL clustering resolved the network into three biochemical modules. The first module corresponded to lipid catabolism and fatty acid β-oxidation, with enrichment of “fatty acid metabolic process” and the KEGG pathway “fatty acid degradation” (**Figure. 3b-c, Supplementary Figure. 7**). Subcellular localization mapped these reactions to peroxisomes and the mitochondrial fatty acid β-oxidation multienzyme complex, indicating dual-compartment lipid utilization as a primary metabolic function of the abdomen. A second module represented ubiquitin-proteasome system (UPS)-mediated proteostasis. Proteasome components constituted the central hubs, with Pros26.4 as the most connected node (8/32, average node degree=2.35). Enriched pathways reflected “proteasome-mediated degradation” and Reactome-level regulation of key signaling substrates, including polyamine, several cell-cycle regulators (e.g. Cyclin D, ODC), and WNT/Hedgehog components (e.g. axin). Multiple subunits of the 26S proteasome (including β4/β7 catalytic core components, Rpt1 ATPase, Rpn11 deubiquitinase, and UBFD1 shuttle factors) formed a highly connected subnetwork.

**Table 3.** Summary of PPI network metrics across tissues.

| Metric | PPI Network Overview |  |  |
| --- | --- | --- | --- |
|  | Head (H) | Gut (G) | Abdomen (A) |
| Nodes | 36 | 149 | 45 |
| Edges | 11 | 181 | 52 |
| Avg node degree | 0.611 | 2.43 | 2.31 |
| Avg local clustering coeff. | 0.315 | 0.345 | 0.572 |
| Expected edges (random) | 3 | 140 | 14 |
| PPI enrichment p-value | 9.77x10<sup>-4</sup> | 4.97x10<sup>-4</sup> | 7.44x10<sup>-15</sup> |

The third module involved nucleotide and redox metabolism. The most connected node was a predicted trifunctional purine biosynthetic protein (9/32), alongside glutamine amidotransferase and AIR carboxylase domain-containing proteins involved in de novo purine biosynthesis. Enrichment of "purine ribonucleoside monophosphate metabolic process", "nucleotide biosynthesis" and "one-carbon pool by folate" indicated coordinated supply of nucleotide precursors and reducing equivalents required to sustain lipid oxidation and ATP-dependent process, such as Rpt1-driven proteasomal substrate unfolding. Additionally, the mitochondrial enzyme AK3 (GTP:AMP phosphotransferase) and pentose phosphate pathway components, including transaldolase, were involved in the subnetwork. These modules collectively suggested a metabolically integrated system in which lipid β-oxidation supplied energy (acetyl-CoA and NADH), nucleotide metabolism supported biosynthetic and redox demands (NADPH via the pentose phosphate pathway), and the proteasome dynamically remodeled the proteome (**Supplementary Table 2**).

### Gut exhibited coordinated metabolic, cytoskeletal, and proteostatic functions

The gut-specific PPI network showed moderate interaction enrichment (PPI enrichment p = 4.97×10^-4^, clustering coefficient 0.345; **Table 3**). MCL clustering of validated gut-specific proteins identified four functional modules (**Figure. 3b-c, Supplementary Figure.76**). The first module was associated with central carbon metabolism and mitochondrial energy production. Hub proteins included pyruvate dehydrogenase E1 subunit α, citrate synthase, aconitate hydratase, and ATP5b. The module was enriched for KEGG ("Citrate cycle", "Glycolysis/Gluconeogenesis") and Reactome pathways ("TCA cycle and respiratory electron transport"). Additionally, the presence of UTP-glucose-1-phosphate uridylyltransferase (UDP-glucose synthesis) indicated linkage between central carbon metabolism and glycoconjugate and glycogen biosynthesis. The second module encompassed protein homeostasis and ubiquitin-mediated regulatory pathways. Hub proteins included CSN8, Rps27A, and UBL4A. Cullin-4A, a scaffold for CRL4 E3 ubiquitin ligases, indicated the presence of targeted protein degradation. Prefoldin that facilitated proper folding of actin and tubulin monomers was also reproduced, thereby linking proteostasis to cytoskeletal integrity. In addition, ER metallopeptidase and cytosolic aminopeptidase were observed, consistent with active proteolytic processing in gut tissue. Notably, enrichment of global genome nucleotide excision repair (GG-NER) pathways suggested that these ubiquitin-dependent regulatory processes may extend beyond proteostasis to contribute to genome maintenance.

Additional proteins formed less interconnected clusters but showed significant enrichment for specific functions. The third module involved intracellular transport, represented by Rab-14, golgin family members, VAMP-B, derlin-2, and TIM13. Functional enrichment analysis highlighted “macromolecule localization”, “protein transport”, “clathrin-mediated endocytosis”, and “endomembrane system”. Those were consistent with active vesicle trafficking and endosomal recycling essential for the gut epithelium’s secretory and absorptive activity. The fourth module captured cytoskeletal organization, with enrichment in “actin cytoskeleton” and “actin filament”. Hub and structural proteins comprised actin, tropomyosin, troponin, myosin regulatory light chain 2, filamin-A, as well as titin and obscurin. Regulatory components of actin dynamics were also detected, including actin-interacting protein 1 (AIP1), which facilitated cofilin-mediated actin disassembly and enabled actin remodeling. α-catenin that coupled adherens junctions to the actin cytoskeleton was also identified for both years. (**Supplementary Table 3**)

### Head was characterized by specialized amino-sugar metabolic and vesicle-mediated transport

The head-specific PPI network (PPI enrichment p = 9.77×10^-4^, clustering coefficient 0.315; **Table 3**) revealed two local functional modules. The first involved amino-sugar catabolism and redox-related processes. A close interaction was observed between GlcNAc-6P deacetylase and a predicted glucosamine-6-phosphate isomerase, enriched for "pentose and glucuronate interconversions" and "small molecule catabolic process" (**Figure. 3b-c, Supplementary Figure. 7**). The deacetylase converted GlcNAc-6P into GlcN-6P, while the isomerase subsequently converted to fructose-6-phosphate, feeding directly into glycolysis^42^. Several linked oxidoreductases were also identified, including aldose reductase, prostaglandin reductase, and trans-1,2-dihydrobenzene-1,2-diol dehydrogenase (**Supplementary Table 4**). These enzymes were associated with redox metabolism and detoxification of reactive aldehydes and lipid peroxidation products.

The second module involved vesicle trafficking and membrane quality control. A representative protein, vesicle-fusing ATPase (the honey bee ortholog of VCP/p97), was identified. This protein had been implicated in (i) membrane dynamics, including homotypic ER and Golgi membrane fusion, and (ii) ubiquitin-dependent protein quality control, particularly the extraction of misfolded proteins from the ER membrane for Endoplasmic Reticulum-Associated Degradation (ERAD)^43,44^. The cellular component term “caveolae” was significantly enriched among head-specific proteins, and also indicated the involvement of membrane microdomains in endocytosis and membrane organization. Co-existence of both modules suggested a coordinated investment in preventing and clearing oxidative protein damage, which may be critical in postLmitotic neurons that cannot dilute damaged proteins through cell division.

### Environment-proteome interactions reveal tissue-specific and reproducible stressor-associated proteomic signatures

To identify dominant environmental influences on tissue proteomes in each year, we performed random-effects meta-analysis on environmentally responsive proteins across crop contrasts. Rather than considering all DEPs, this approach focused on proteins showing consistent patterns of expression transition across environments, including both conserved expression blocks (no significant crop differences) and variable blocks containing at least one significant crop contrast. By emphasizing the configuration of expression changes across contrasts rather than individual pairwise effect sizes, the analysis captured coordinated environmental response patterns shared across crops. Consistent with this interpretation, the resulting environment-sensitive proteins exhibited moderate but systematic expression shifts, with mean and maximum effect sizes ranging from 0.22-0.27 and 0.44-0.48 log_2_FC, respectively.

These environment-associated effects were tissue-specific and changed between years. In 2020, CRA exerted the strongest upregulatory influence on abdomen-dominant proteins (pooled effect μ = +0.152), indicating their strongest overall increase in expression relative to other crops. CAS dominated the gut responses (μ = +0.168), whereas HBB had the largest absolute effect in the head (μ = -0.175). In 2021, the pattern shifted: CRA imposed the largest suppressive effect on both head (μ = - 0.325) and gut (μ = -0.370), while CAS exerted the primary upregulating effect on abdomen (μ = +0.261; **Figure. 4a-b, Supplementary Table 5**).

**Figure 4.**
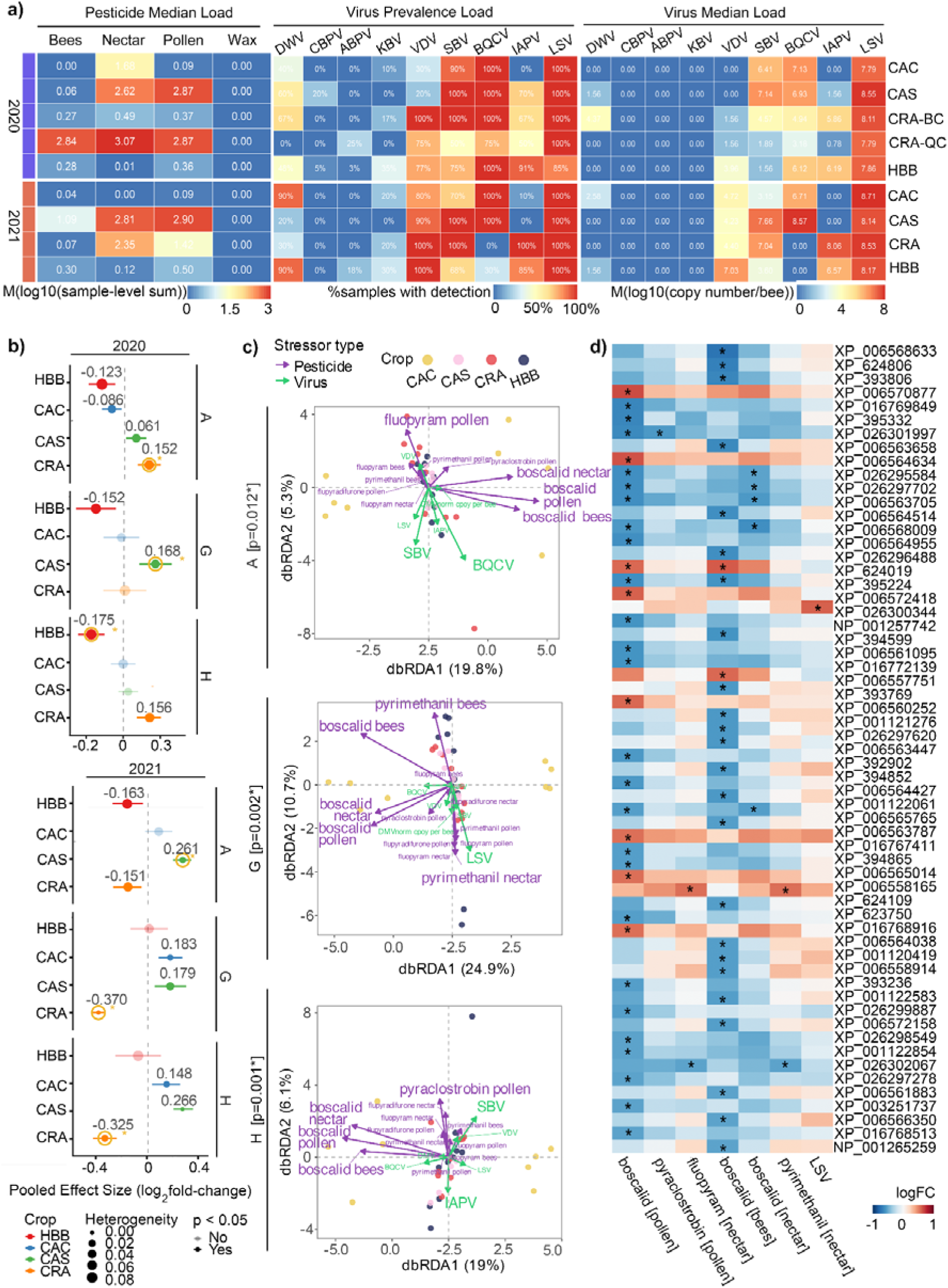
Environment-proteome interactions in honey bee head (H), abdomen (A), and gut (G) tissues reveal reproducible stressor-associated proteomic responses across four crops (CAC = commodity canola; CAS = seed canola; CRA = cranberry; HBB = highbush blueberry) and two years (2020-2021). (**a**) Pesticide exposure (median load across bee, nectar, pollen, and wax matrices) and virus prevalence and median load across crops and years. Values represent normalized exposure estimates as described in MATERIALS and METHODS. M refers to the median in the color scale bar. **(b)** Meta-analysis of proteins exhibiting crop-to-crop transitions across years. Effect sizes represent pooled log-fold changes across comparisons; significance thresholds are indicated in MATERIALS and METHODS. **(c)** Distance-based redundancy analysis (dbRDA) of tissue-specific proteomic profiles constrained by environmental stressors in 2020. Axes represent constrained ordination of proteomic variation associated with stressor gradients. **(d)** Heatmap of significant protein-stressor correlations among proteins pre-filtered by envfit in 2020 (envfit p < 0.05; Spearman FDR < 0.05).

The most prominent temporal pattern was a reversal of CRA-associated effects between years, from a positive abdomen effect in 2020 (+0.152) to negative effects across tissues in 2021, most strongly in the head (-0.325) and gut (-0.370). This temporal shift occurred against a changed CRA sampling context: the 2020 CRA group included both BC and QC sites, whereas the 2021 CRA group represented BC sites only. Within the BC CRA cohort, this was accompanied by an increase in IAPV prevalence and raw median viral copy number, while measured pesticide residues differed between the CRA site-year groups (**Figure. 4a**, **Supplementary Tables 6-7**). Because crop system, geography, beekeeper management, surrounding agricultural landscape and mid-bloom sampling date are partially confounded in this field design, we interpret these patterns as crop ecosystem/year-associated environmental signatures rather than isolated crop effects. Consistent with this interpretation, CAS showed consistently high measured pesticide residue loads, particularly in nectar and pollen, whereas CAC and HBB showed lower residue burdens in this dataset; these differences likely reflect both crop system and surrounding landscape or management context.

Beyond the direction of the pooled effect, the meta-analysis also distinguished whether proteins within each tissue responded coherently or heterogeneously. The magnitude of between-protein heterogeneity (τ^2^) and the proportion of variance attributable to true effect-size differences (I^2^) varied substantially across tissue-crop-year combinations. CRA in gut 2021 showed near-zero heterogeneity (τ^2^ = 0; I^2^ = 0.1%). This indicated that gut-dominant proteins responded with highly concordant effect sizes and in a consistent suppressive direction. Functionally, the suppressed protein set was enriched for biosynthetic and energy-demanding processes related to translation, mitochondrial energy metabolism, and proteasome-mediated protein turnover. Representative proteins included NOP56, 40S ribosomal protein SA (RPSA), ATP synthase α, and proteasome subunit β4 (**Supplementary Table 8**).

In contrast, CRA in abdomen 2020 exhibited substantial heterogeneity (τ^2^ = 0.036-0.089; I^2^ up to 90%), reflecting pronounced variability in either magnitude or direction of effect sizes across proteins. This responses spanned diverse functional programs within the fat body, including membrane glycoprotein quality control (FUT8), translation and RNA metabolism (eIF4E-type 2, PABP, U4/U6 Prp3, Tudor domain-containing protein 7, and ATPase ASNA1), as well as a broader set of metabolic and cellular maintenance processes supporting one-carbon metabolism (ALDH1L1), TCA cycle substrate transport (mitochondrial dicarboxylate carrier), phospholipid biosynthesis (AGPAT), DNA replication and mitochondrial function (POLDIP2), and stress-responsive phosphatase regulation (PP5).

### Tissue***-***specific functional properties under dominant crops

Head responses were suppressive in both years, shifting from an HBB-associated effect in 2020 (μ = -0.175) to a stronger CRA-associated effect in 2021 (μ = -0.325). In the head samples, the dominant crop shifted from HBB in 2020 to CRA in 2021 , with both conditions associated with suppressive effects but differing in magnitude and heterogeneity. The HBB-associated suppression in 2020 involved proteins related to neuronal biosynthetic and metabolic functions, including translation (e.g., ribosomal proteins L14 and L23), mitochondrial dynamics and energy metabolism (mitofusin/Marf, V-type proton ATPase), and neuroendocrine processing (neuroendocrine convertase 2). Additional proteins related to synaptic organization and vesicle dynamics (e.g., gephyrin, Rab-35) were also affected. In contrast, CRA-associated suppression in 2021 involved proteins linked to transcriptional regulation and membrane trafficking (RPB2, splicing factor 45, and AP-2 complex subunit α), as well as components of ionic homeostasis such as Na^+^/K^+^-ATPase α (**Supplementary Table 8**).

The gut showed the strongest functional switch, from CAS-associated induction in 2020 (μ = +0.168) to CRA-associated suppression in 2021 (μ = -0.370). The CAS-associated upregulation in 2020 primarily involved proteins supporting epithelial defense and metabolic activity, including membrane trafficking and secretion (e.g., Golgin-97, HOOK3), mitochondrial function (OXA1L, DHX30), and proteostasis (PDIA6, HERC4). Detoxification-related components were also represented, notably PAPSS, which supplied activated sulfate for phase II conjugation reactions. Together, these proteins indicated an active, protective response of the gut epithelium to chemical exposure. In contrast, CRA-associated suppression in 2021 involved a coordinated reduction in core cellular functions, particularly those related to translation, energy metabolism, and protein turnover, as described above (**Supplementary Table 8**).

Abdomen responses were instead characterized by positive effects in high-residue crop-system contexts. In abdomen tissue, the dominant upregulatory crop shifted from CRA in 2020 (μ = +0.152) to CAS in 2021 (μ = +0.261), with both conditions carrying high pesticide load. In 2020, CRALassociated proteins were enriched for a broad homeostatic tuning of biosynthetic and metabolic processes described above. CAS-associated upregulation in 2021 exhibited a more coordinated biosynthetic response involved HBBLstable proteins (eIF5, sepiapterin reductase) and CACLstable proteins (MCT3, nucleolin, ILF2, CK2β, CD2AP) that supported protein synthesis, metabolic flux, and cellular remodeling under sustained pesticide exposure (**Supplementary Table 8**).

Next, we tested the reproducibility of the predictive power of proteins identified as crop-specific and tissue-specific. We took all proteins identified as tissue-specific from 2020 and 2021, trained a logistic regression model and a random forest model on the 2020 protein data, and then tested them on the 2021 data. We found that the Random forest model performed better than logistic regression (Accuracy = 0.99 and 0.75 respectively, **Supplementary Figure 8i**), and that only two proteins were required to pinpoint the exact tissue to which a sample belonged (**Supplementary Figure 9, 10**). We performed a similar analysis for proteins identified as crop-specific, but the model lacked predictive power.

### The Gut as the Principal Site of Protein-Boscalid Crosstalk

We then asked whether measured pesticides and viral loads could explain the crop ecosystem/year-associated proteomic structure. Eighteen stressor variables were included as predictors, comprising 4 pesticide residues each measured across pollen, nectar, and bee tissues, and six RNA viruses (DWV, VDV, SBV, BQCV, IAPV, LSV). Pesticide exposure was expected to remodel the tissue proteome, potentially shifting both the central tendency of protein composition (a location effect) and the variability among samples (a dispersion effect), as chronic agrochemical stress could either drive directional change or destabilize proteomic homeostasis. To test this, we partitioned variation in protein composition using PERMANOVA with marginal effects (adonis2, Bray-Curtis dissimilarity) and assessed multivariate homogeneity of dispersion using betadisper, analysing each year separately. Exposure explained a negligible and non-significant fraction of compositional variation in both years (**Supplementary Figure 11, Table 4**).

**Table 4.** PERMANOVA and betadisper results for pesticide exposure effect on protein composition.

| Year | Test | F-stat | R<sup>2</sup> | p-value |
| --- | --- | --- | --- | --- |
| 2020 | PERMANOVA (exposure, margin) | 0.930 | 0.002 | 0.472 |
|  | betadisper (exposure) | 0.203 | NA | 0.637 |
| 2021 | PERMANOVA (exposure, margin) | 1.878 | 0.002 | 0.118 |
|  | betadisper (exposure) | 0.101 | NA | 0.766 |

However, this global pattern masked substantial heterogeneity among individual stressors. Two stressors showed strong cropLrelated stratification, with >60% of their variance explained by crop identity (ICC > 0.6): pyrimethanil and IAPV. The remaining stressors showed lower crop association, with ICC values ranging from 0.083 to 0.588. In 2020, honey bee proteomic variation was predominantly shaped by pesticide exposure across all tissues after controlling for crop ecosystem effects, with the gut showing the strongest overall responsiveness. The global db-RDA models were statistically significant for abdomen (p = 0.016, R^2^ = 55.9%), gut (p = 0.002, R^2^ = 60.8%), and head (p = 0.001, R^2^ = 58.7%). Therefore, more than half of the total proteomic variation could be explained by the measured stressor panel. In all tissues, the first constrained axis (CAP1; 19-25%) was primarily aligned with pesticide exposure rather than viral load (**Figure. 4c**, **Table 5**).

**Table 5.** Global db-RDA ANOVA on the tissue-specific protein.

| Year | Tissue | n | F | p-value | R<sup>2</sup> (%) | CAP1 (%) | CAP2 (%) |
| --- | --- | --- | --- | --- | --- | --- | --- |
| 2020 | Abdomen | 38 | 1.336 | 0.016 | 55.86 | 19.8 | 5.3 |
| 2020 | Gut | 38 | 1.639 | 0.002 | 60.83 | 24.9 | 10.7 |
| 2020 | Head | 38 | 1.503 | 0.001 | 58.74 | 19.0 | 6.1 |

At the level of individual stressors, boscalid emerged as the only consistently significant driver across tissues (**Supplementary Table 9**). In the abdomen and gut, boscalid in pollen and nectar showed significant marginal-ANOVA associations with proteomic variation (p ≤ 0.01), while in the head all three exposure routes were significant (p = 0.002-0.039). Viral variables were largely non-significant, except SBV in the head (p = 0.041). Across all three tissues, boscalid in pollen consistently ranked as the top-contributing stressor by effect size (variance: A = 27.23, G = 24.47, H = 23.60), with boscalid in nectar second (variance: A = 17.69, G = 21.07, H = 18.43).

We next examined whether these stressor-level patterns were reflected by individual protein-stressor associations. The gut was the principal site of protein-stressor crosstalk, accounting for 60 unique proteins across seven distinct stressors (**Figure. 4d, Supplementary Table 10**). Of all proteins with significant FDR (asterisks), the majority interacted with the boscalid in pollen and in bees, and almost all of these flagged correlations were negative (cool blue). Therefore, the response between gut proteins to boscalid exposure was monotonically negative. This pattern was corroborated by the regression-based validation in 2021. 27 protein-stressor pairs reproduced across years (**Supplementary Figure. 12, Supplementary Table 11**), and the implicated proteins converged onto a limited number of functional modules, including mitochondrial energy metabolism, protein quality control, vesicle trafficking, and gut-specific carbon and biosynthetic pathways.

### Multi-omics integration preserves gut proteomic stressor programs and links them to transcriptomic and microbiome variation

The single-proteomics MOFA and db-RDA analyses above identified boscalid as the primary chemical driver of gut proteomic variation and revealed tissue-specific stressor response programs. To assess whether these proteomic patterns were robust across measurement platforms and to evaluate whether protein-level variation coordinated with transcriptomic and microbiome changes, we performed an integrated MOFA2 analysis jointly modelling proteomic (4068 proteins), transcriptomic (5000 genes), and microbiome (48 taxa) data from gut tissue. Single-omics MOFA models trained independently on each platform revealed that proteomics consistently captured more total variance and yielded more significant factor-stressor associations than either transcriptomics or the microbiome alone (**Figure. 5a-c**). In factor-phenotype association comparisons, the multi-omics model matched or exceeded the single-omic models for 5 of 12 covariates with at least one significant hit (**Figure. 5d**). Notably, the multi-omics model did not diminish the detection of key stressor associations but instead placed them within a broader biological context by linking protein-level responses to coordinated transcriptomic Program 2 (**Figure. 5f-g**). Among the highest-loading microbial features, *Gilliamella apicola* and *Gilliamella* sp. A7, both core members of the honey bee gut symbiont community, varied along the same latent axis as the proteomic and transcriptomic features. These results suggest that the CAS-associated pathogen axis was not restricted to protein abundance, but reflected coordinated variation across host molecular state and gut microbial composition.

**Figure 5.**
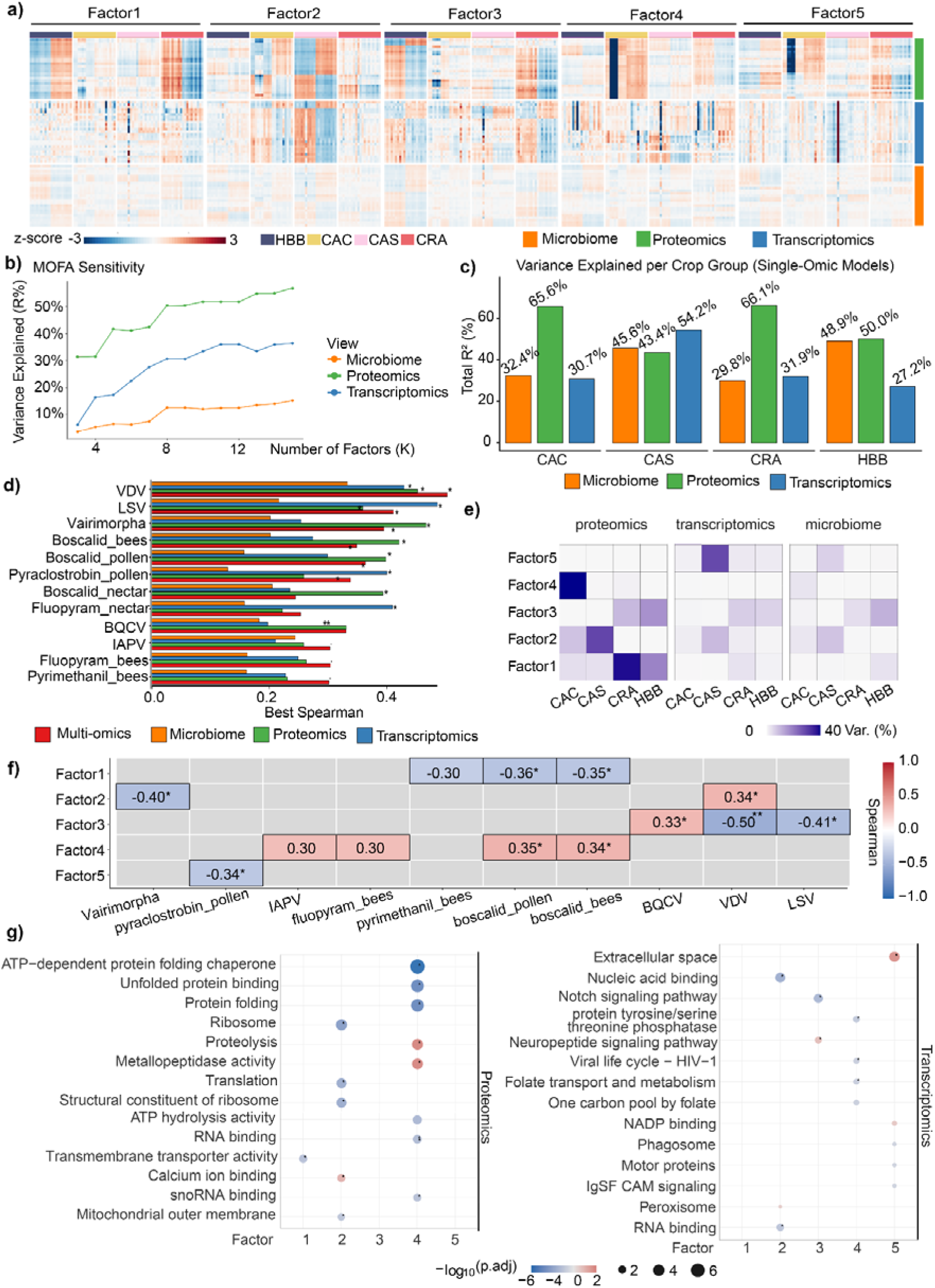
Multi-omics factor analysis identifies coordinated microbiome, proteomic, and transcriptomic programs in the gut. (**a**) Z-scored protein abundance across samples within each tissue. Colors represent standardized protein abundance (z-scores) computed across samples for each protein within each tissue. Red indicates higher-than-average abundance and blue indicates lower-than-average abundance relative to the protein-specific mean. Each row represents a protein grouped by omics layers. Columns represent samples ordered by crop group (CAC = commodity canola; CAS = seed canola; CRA = cranberry; HBB = highbush blueberry). **(b)** Sensitivity analysis of MOFA models across latent factor numbers (K = 3-15). Total variance explained (R^2^) is shown for each omics view. The optimal model was selected at K = 9 factors. **(c)** Variance explained (R^2^) by single-omics MOFA models across crop groups for microbiome, proteomic, and transcriptomic datasets. **(d)** Comparison of factor-phenotype associations across single-omics and multi-omics models based on maximum Spearman correlation coefficients. Significant associations are indicated by *FDR < 0.05. **(e)** Variance explained by each latent factor across omics views and crop groups. **(f)** Associations between latent factors and environmental stressors. Significant associations are indicated by *FDR < 0.05 and **FDR < 0.01. **(g)** Gene Ontology (GO) enrichment analysis of high-loading features associated with latent factors in the proteomic and transcriptomic views. Dot size represents gene count and colour indicates enrichment significance.

The remaining integrated axes separated additional virus- and pesticide-linked gut states. One axis connected all three molecular layers primarily in CRA and HBB and was associated with VDV (p=-0.504), LSV (p=-0.412) and BQCV (p=+0.331; **Figure. 5e-g**). Its leading microbiome features included *Bifidobacterium actinocoloniiforme* and *Bifidobacterium asteroides*, obligate gut symbionts involved in carbohydrate fermentation and pathogen resistance, indicating that virus-associated host variation was accompanied by shifts in functionally relevant gut bacteria. A separate CAC-active axis explained 49.2% of proteomic variance and recapitulated the pesticide-enriched profile of proteomic Program 4, with significant associations for boscalid in pollen (p=+0.354), boscalid in bees (p=+0.345), fluopyram in bees (p=+0.304) and IAPV (p=+0.304) at FDR<0.1 (**Figure. 5e-g**).

Together, these results show that multi-omics integration preserved the principal gut proteomic stressor programs while linking them to coordinated transcriptomic and microbiome variation.

### Temporal validation of compact XGBoost+RFECV crop-ecosystem-associated molecular signatures

After the descriptive and multi-omics analyses identified crop-associated molecular programs across tissues and in the gut, we next asked whether these patterns contained a compact component that could be transferred across years. This final analysis retained an explicitly supervised machine-learning component: LIMMA-derived crop-contrast features were used as broad descriptive references, whereas XGBoost screening, RFECV selection, and Random Forest validation were used to evaluate whether sparse tissue-specific signatures selected in 2020 remained predictive in 2021^33,45–47^.

To preserve the independence of the 2021 validation cohort, all supervised screening, feature selection, model optimisation, and model fitting were restricted to 2020 samples. The resulting feature panels were then fixed and applied unchanged to the independent 2021 cohort. The workflow is summarised in **Figure. 6a**: LIMMA provided the descriptive reference for crop-associated structure, XGBoost reduced the candidate feature space, RFECV selected sparse panels, and Random Forest classifiers tested temporal transfer of the selected signatures.

**Figure 6.**
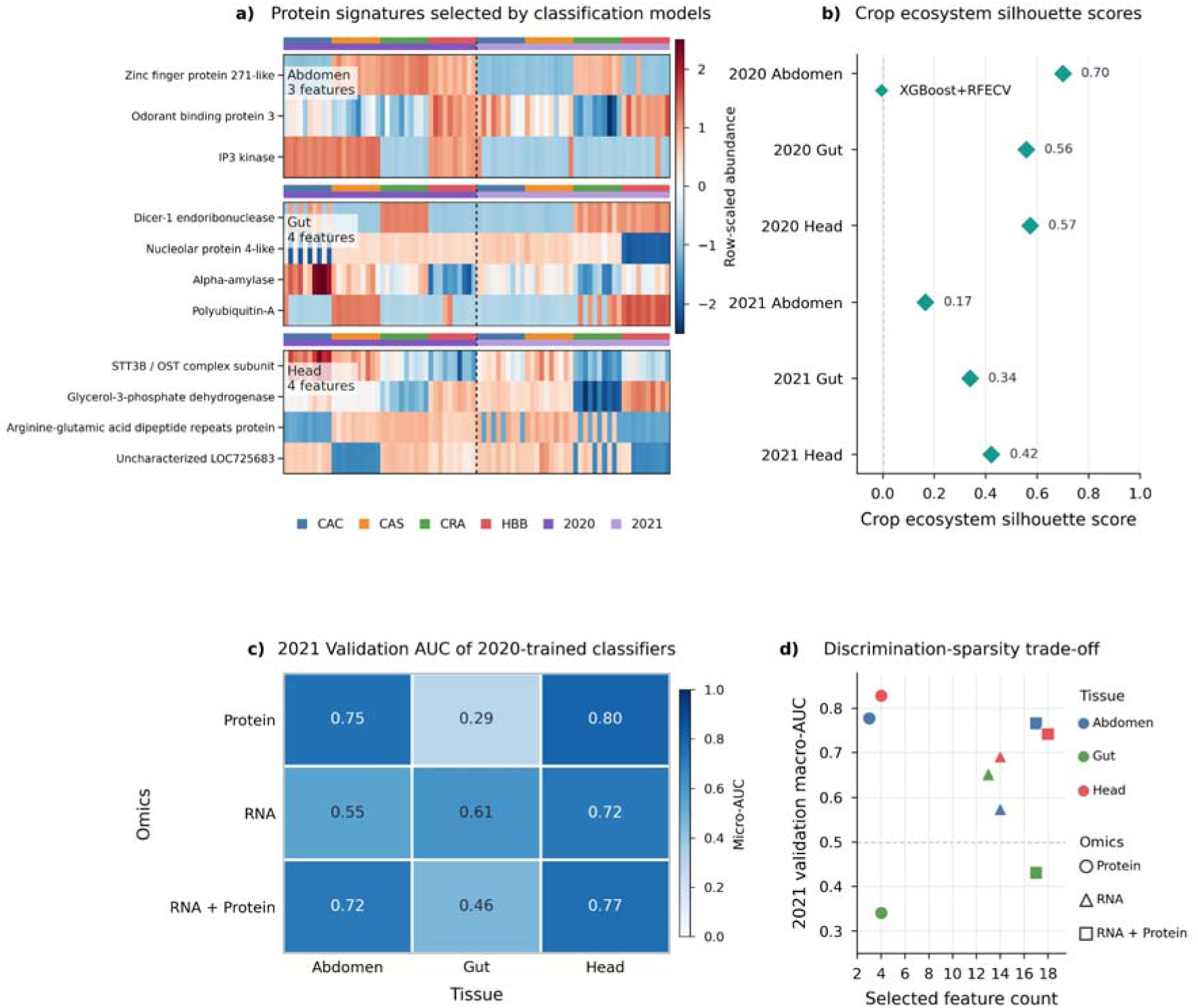
Temporal validation of compact crop-ecosystem-associated molecular signatures. **a)** Protein signatures selected by 2020-trained classification models. Row-scaled abundance heatmap of compact XGBoost+RFECV-selected protein signatures for abdomen, gut and head. Rows represent selected proteins and columns represent samples ordered by crop ecosystem and year. Top annotations indicate crop ecosystem and sampling year; the dashed vertical line separates the 2020 training cohort from the independent 2021 validation cohort. **b)** Crop ecosystem separation of selected protein signatures. Crop ecosystem silhouette scores calculated from the selected protein-signature embeddings for each tissue and year. Higher values indicate stronger separation of samples by crop ecosystem. **c)** Independent-year validation of 2020-trained classifiers. Micro-average AUC values for Random Forest classifiers trained and tuned using 2020 data and evaluated without further feature selection in the 2021 cohort. Rows indicate omics input type and columns indicate tissue. **d)** Discrimination-sparsity trade-off. Relationship between selected feature count and 2021 validation macro-average AUC across tissue and omics settings. Point colour indicates tissue and point shape indicates omics input type.

The XGBoost+RFECV workflow selected highly compact protein signatures for each tissue, consisting of three abdomen proteins, four gut proteins, and four head proteins **(Figure. 6b)**. Despite this strong reduction in dimensionality, the row-scaled heatmap retained visible abundance structure across samples ordered by year, crop, and mapped location. Abdomen and head showed the clearest compact protein-associated structure, whereas the gut displayed a more heterogeneous pattern. A direct heatmap comparison between the broad LIMMA-derived descriptive reference and the compact supervised protein panel is provided in the supplementary material **(Supplementary Figure. 13)**. We next quantified whether the LIMMA reference features and the fixed 2020-derived supervised panels separated samples more strongly by crop than by mapped geographic location. For each UMAP embedding, we calculated the difference between crop and location silhouette scores^31,39^:

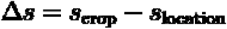

Positive values indicate stronger separation by crop than by mapped location. In the training year, the compact supervised panels preserved crop-associated structure in abdomen and gut despite using far fewer features than the LIMMA references, whereas the LIMMA reference retained stronger separation in head **(Figure. 6c)**. In 2021, the fixed abdomen panel remained similar to the LIMMA descriptive reference, while LIMMA showed stronger crop-over-location separation in gut and head. The tissue-year-separated UMAP embeddings underlying this summary are shown in the supplementary material **(Supplementary Figure. 14)**. Together, these patterns indicate that the sparse predictive panels did not simply reproduce the broad LIMMA structure, but instead captured a smaller supervised component of the crop-associated signal whose transferability varied by tissue.

Independent-year prediction further showed that temporal transfer was tissue and modality dependent **(Figure. 6d)**. Protein-only panels supported strong 2021 discrimination in abdomen and head, with micro-average AUC values of 0.749 and 0.799, respectively. In contrast, protein-based prediction was weaker in the gut: the protein-only gut model reached a micro-average AUC of 0.287, whereas RNA provided the strongest gut performance with a micro-average AUC of 0.612. Early RNA + protein fusion did not uniformly improve temporal validation performance, reaching a micro-average AUC of 0.716 in abdomen and 0.765 in head but remaining below the best single-omic model in these tissues and below RNA alone in gut. Detailed validation metrics, including precision, recall, F1-score, macro-average AUC, micro-average AUC, and selected feature counts, are provided in the supplementary material **(Supplementary Figure. 15)**.

To evaluate whether stronger discrimination required larger feature panels, we compared 2021 macro-average AUC against selected feature count (Figure. 6e). Macro-average AUC was calculated as^40^:

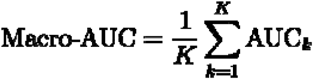

where K is the number of crop classes and AUC_k_ is the one-vs-rest AUC for crop class k. The strongest discrimination-sparsity trade-offs were observed for the protein-only abdomen and head models. The abdomen protein model achieved a macro-average AUC of 0.778 using only three features, and the head protein model achieved a macro-average AUC of 0.829 using only four features. Larger RNA and RNA + protein panels did not guarantee higher validation performance, indicating that independent-year discrimination was not simply a function of feature-set size.

Together, these results extend the descriptive and multi-omics analyses by testing whether crop-associated molecular structure could be compressed into temporally transferable signatures. LIMMA-derived features provided a broad year-specific reference for crop-associated structure, while XGBoost+RFECV compressed part of this structure into sparse panels selected from 2020 only.

These fixed panels supported independent-year prediction most strongly in abdomen and head, whereas gut remained a weaker and more heterogeneous predictive context. Thus, crop-associated molecular signals showed tissue-dependent temporal transferability, and compact supervised feature panels provided interpretable predictive summaries without requiring broad univariate feature sets.

## Conclusions

Tissue-resolved proteomics separated two layers of molecular variation in field-exposed honey bees. The first was a stable tissue architecture: abdomen, gut, and head retained distinct proteomic programs across crop ecosystems and years. The second was an environmentally structured layer, in which crop-ecosystem and year effects selectively remodelled these tissue programs. This organization demonstrated that tissue identity remained the dominant source of reproducible proteomic structure, while specific field contexts still produced large, tissue-dependent shifts.

The strongest environmental contrast was the year-dependent reversal in CRA, from abdomen-dominant induction in 2020 to broad gut and head suppression in 2021. CAS showed a different pattern, with upregulation of gut proteins linked to epithelial maintenance and defense under the highest pesticide-load context. These patterns argue against a single generalized stress response. Instead, they indicate that field exposures produce distinct tissue-level response modes, shaped by the combination of crop ecosystem, year, pathogen burden, and pesticide profile.

The gut emerged as the main site where proteomic, chemical, and cross-omic signals converged. Boscalid was the most reproducible chemical correlate of proteomic variation after accounting for crop-ecosystem-level structure. Of 27 reproducible protein-stressor pairs, 26 were negative associations between gut-specific proteins and boscalid measured in pollen or bee tissue (**Supplementary Figure 12, Supplementary Table 11**). These associations involved gut proteins linked to energy-dependent processes, including mitochondrial metabolism, vesicle trafficking, nutrient transport, and biosynthesis, consistent with boscalid inhibition of succinate dehydrogenase and reported effects on honey bee mitochondrial respiration^48^. Integrated gut multi-omics preserved the main proteomic stressor programs and linked them to coordinated transcriptomic and microbiome variation, indicating that the pesticide- and pathogen-associated signatures were not restricted to protein abundance alone. The temporal validation analysis further showed that some crop-ecosystem-associated structure could be compressed into compact protein panels with independent-year transferability, particularly in abdomen and head, whereas gut variation remained more heterogeneous and context dependent.

These results should be interpreted as field associations rather than causal effects. The design intentionally captured realistic agroecosystem variation, but crop ecosystem, region, beekeeper identity, sampling composition, and stressor burden were not fully orthogonal. Future controlled exposure studies will be needed to test whether boscalid and IAPV directly drive the proteomic changes observed here, alone or in combination. More balanced field sampling, denser longitudinal designs, and spatial or single-cell proteomic approaches^49^ would further resolve whether bulk tissue programs arise from broad tissue remodeling or from shifts in specific cell populations, such as fat-body trophocytes and oenocytes^50^.

Overall, this study shows that honey bee proteomes in crop agroecosystems are organized by stable tissue identity but remain sensitive to environmentally structured chemical and pathogen pressures. Gut-resolved proteomics, in particular, provides a sensitive readout of sublethal pesticide-associated remodeling under field conditions and offers a tractable entry point for mechanistic validation of chronic stressor effects in managed pollinators^51^.

## Supporting Information

**Supplementary Table 1. Tables with samples and stressor factors** 969 **Supplementary Table 2.**
|  |  |
| --- | --- |
| sup-0001-Data1 | Supplementary Table 1 |
| sup-0002-Data2 | Supplementary Tables 2-11 |
| sup-0003-Data3 | Supplementary Figures 1-15 |

**Supplementary Table 1. Tables with samples and stressor factors**

**Supplementary Table 2. List of abdomen-specific protein names and their accessions**

**Supplementary Table 3. List of gut-specific protein names and their accessions**

**Supplementary Table 4. List of head-specific protein names and their accessions**

**Supplementary Table 5. Random Effect Meta-analysis for crop-dominant effects across tissues and years**

**Supplementary Table 6. Environmental burden profiles (pesticides) per crop and year**

**Supplementary Table 7. Environmental burden profiles (virus) per crop and year**

**Supplementary Table 8. Environmentally sensitive proteins**

**Supplementary Table 9. Significant stressors (per-term ANOVA)**

**Supplementary Table 10. Spearman correlations passing FDR < 0.05 in 2020 .**

**Supplementary Table 11. Proteins of different origins showing reproducible negative association with boscalid across years in the gut or head**

**Supplementary Figure 1. Crop structure and dispersion in proteomic space**

**Supplementary Figure 2. Near/far exposure sensitivity analysis in the global protein expression space.**

**Supplementary Figure 3. Gene Ontology (GO) enrichment analysis of proteins with high loadings in tissue-specific latent programs.**

**Supplementary Figure 4. Heatmap showing (a) GO and (b) KEGG enrichment patterns of tissue-specific DEPs across tissue pairwise contrasts.**

**Supplementary Figure 5. Heatmap showing scaled expression patterns (logFC) of tissue-specific proteins across tissue contrasts (A vs G, A vs H, and G vs H) in different crops and years.**

**Supplementary Figure 6. Global PPI networks constructed from tissue-specific DEPs using STRING analysis.**

**Supplementary Figure 7. Functional enrichment bubble plots of tissue-specific PPI subnetworks in abdomen, gut, and head tissues.**

**Supplementary Figure 8. Confusion matrices showing bee tissue predictions for the 2021 dataset using a random forest (i) and logistic regression (ii) models trained on the 2020 dataset.**

**Supplementary Figure 9. Decision tree, built based on top random forest features**

**Supplementary Figure 10. Bar plot shows proteins selected by multinomial LASSO logistic regression for bee tissue classification.**

**Supplementary Figure 11. PERMANOVA (i) and betadisper (ii) results for pesticide exposure effect on protein composition**

**Supplementary Figure 12. Representative 2020 protein-stressor association (selected after envfit pre-filtering) independently reproduced in 2021 using sign-consistent FDR-significant validation criteria.**

**Supplementary Figure 13. Heatmap comparing broad LIMMA-derived descriptive features with compact XGBoost+RFECV-selected protein features.**

**Supplementary Figure 14. Tissue-year-separated UMAP visualisation of LIMMA-derived descriptive reference features and fixed XGBoost+RFECV protein panels.**

**Supplementary Figure 15. Detailed independent-year prediction metrics for XGBoost+RFECV-selected feature panels.**

## Data and Code Availability Statement

The mass spectrometry data have been deposited in the ProteomeXchange Consortium via MassIVE (Mass Spectrometry Interactive Virtual Environment) partner repository under the dataset identifier PXD062819 for HBB and CRA, and via PRIDE (PRoteomics IDEntifications Database) partner repository under the dataset identifier PXD078424 for CAS and CAC. The transcriptome data are available upon request by email to the authors. The raw metagenome sequencing data are available in BioProject PRJNA999720 in the NCBI BioProject database. The code will be released on GitHub upon publication.

## Acknowledgments

This work was funded by the Government of Canada through Agriculture and Agri-Food Canada (AAFC) Genomics Research and Development Initiative (GRDI) funding (AAFC J-002368), Genome Canada (LSARP #16420), and the Ontario Genomics Institute (OGI-185). Mass spectrometry infrastructure used here was supported by the Canada Foundation for Innovation, the BC Knowledge Development Fund, the Life Sciences Institute, and Genome BC (374PRO). This research was enabled in part by support provided by the Digital Research Alliance of Canada (alliancecan.ca) and the AAFC Biocluster. AKR was supported by the HSE University basic research program. We thank Alison McAfee and Yuming Shi for helpful comments on the analysis and Figures. We would like to thank the following assistants, post doc and students for their contributions to this project: Bradford Vinson, Renee Teo, Irene Yu, Inna Kostiuk, Irina Chua, Francesco Babini, Elizabeth Adefolaju, Rhonda Thygesen, Clarice Bryant, Zoe Rempel, Derek Micholson, Carolyn Currie, Kira Peters, Vivane Beger, Abdullah Ibrahim, Kelan Lynch, Diana Tran. We are also grateful to Bradford Vinson, Irene Yu, and Greg Stacey for their technical support. We also acknowledged the landowners, beekeepers and field crews, without whom the field component could not have been completed.

## Authors contribution

**Conceptualization:** Huan Zhong, Peipei Zhong, Amro Zayed, M. Marta Guarna, Stephen F. Pernal, Robert W. Currie, Pierre Giovenazzo, Shelley E. Hoover, and Leonard J. Foster. **Methodology:** Huan Zhong, Peipei Zhong, Junseo Park, Aleksandra Kozlova-Ryabova, , Renata Moravcova, Wendy W. T. Fang, Kyung-Mee Moon, Xiaojing Yuan, Jason C. Rogalski, Ida M. Conflitti, Mateus Pepinelli, Syed Abbas Bukhari, Sarah K. French, Lan Tran, Heather Higo, Julia Common, Jeff Kearns, Amanda S. Gregoris, Elizabeth Walsh, Lance Lansing, Morgan Cunningham, Jonathan Ho, Thomas B. Deckers, Jackie Zorz, Aidan Jamieson, Lynae P Ovinge, Jeff D Kearns and Shelley E. Hoover, Robert W. Currie, Stephen F. Pernal. **Validation:** Huan Zhong, Peipei Zhong, Junseo Park, and Aleksandra Kozlova-Ryabova. **Formal analysis:** Huan Zhong, Peipei Zhong, Junseo Park, and Aleksandra Kozlova-Ryabova. **Data curation:** Huan Zhong, Peipei Zhong, Renata Moravcova, Ida M. Conflitti, Mateus Pepinelli, Syed Abbas Bukhari, Sarah K. French, Lan Tran, Heather Higo, Julia Common, Jeff Kearns, Amanda S. Gregoris, Elizabeth Walsh, Lance Lansing, Morgan Cunningham, Jonathan Ho, Thomas B. Deckers, Jackie Zorz, Aidan Jamieson, Lynae P Ovinge, Jeff D Kearns and Shelley E. Hoover. **Writing - original draft:** Huan Zhong, Peipei Zhong, Junseo Park, Aleksandra Kozlova-Ryabova, and Leonard J. Foster. **Writing - review and editing:** all authors. **Funding acquisition:** Leonard J. Foster, Amro Zayed, Shelley E. Hoover, Stephen F. Pernal, Pierre Giovenazzo, M. Marta Guarna, and Robert W. Currie. **Resources:** Amro Zayed, Leonard J. Foster, Stephen F. Pernal, M. Marta Guarna, Pierre Giovenazzo, Shelley E. Hoover, Robert W. Currie, Rodrigo Ortega Polo, Hosna Jabbari, Huan Zhong. **Project administration:** Amro Zayed, Leonard J. Foster, M. Marta Guarna, Stephen F. Pernal, Shelley E. Hoover, Pierre Giovenazzo, Robert W. Currie, Ida M. Conflitti. **Supervision:** Leonard J. Foster, Amro Zayed, M. Marta Guarna, Stephen F. Pernal, Shelley E. Hoover, Pierre Giovenazzo, Robert W. Currie, Hosna Jabbari, Rodrigo Ortega Polo, and Huan Zhong.

## Abbreviations

CAC: Commodity canola (crop agroecosystem)
CAS: Seed canola; hybrid seed-production canola (crop agroecosystem)
CRA: Cranberry (crop agroecosystem)
HBB: Highbush blueberry (crop agroecosystem)
A: Abdomen (tissue)
G: Gut / midgut (tissue)
H: Head (tissue)
BC: British Columbia
QC: Québec
ABPV: Acute bee paralysis virus
BQCV: Black queen cell virus
CBPV: Chronic bee paralysis virus
DWV: Deformed wing virus
IAPV: Israeli acute paralysis virus
IRES: Internal ribosome entry site
KBV: Kashmir bee virus
LSV: Lake Sinai virus
SBV: Sacbrood virus
VDV: Varroa destructor virus (VDV-1 / Deformed wing virus type B)
BCA: Bicinchoninic acid (protein quantification assay)
C18: Octadecyl-bonded silica (reversed-phase chromatography material)
CAA: Chloroacetamide (cysteine alkylating agent)
DIA-NN: Data-Independent Acquisition by Neural Networks (proteomics search software)
DIA-PASEF: Data-Independent Acquisition-Parallel Accumulation Serial Fragmentation
DTT: Dithiothreitol (reducing agent)
LC-MS/MS: Liquid chromatography-tandem mass spectrometry
LFQ: Label-free quantification (quantitative mass spectrometry)
SDS-PAGE: Sodium dodecyl sulfate-polyacrylamide gel electrophoresis
STAGE-tip: STop-And-Go-Extraction tip (peptide desalting/micro-purification)
Tris-HCl: Tris(hydroxymethyl)aminomethane hydrochloride (buffer)
UHPLC: Ultra-high-performance liquid chromatography
timsTOF: Trapped ion mobility spectrometry-time of flight (mass spectrometer)
ANOVA: Analysis of variance
AUC: Area under the (receiver operating characteristic) curve
CLR: Centered log-ratio (transformation)
CPM: Counts per million (RNA-seq normalisation)
db-RDA (dbRDA): Distance-based redundancy analysis
CAP1 / CAP2: Constrained analysis of principal coordinates, axes 1 and 2
MOFA / MOFA2: Multi-Omics Factor Analysis (version 2)
PCA: Principal component analysis
PC1 / PC2: Principal component 1 / principal component 2
PERMANOVA: Permutational multivariate analysis of variance
PERDISP (betadisper): Permutational analysis of multivariate dispersions
R²: Coefficient of determination (proportion of variance explained)
RFECV: Recursive feature elimination with cross-validation
RandomizedSearc hCV: Randomized search with cross-validation (hyperparameter tuning)
XGBoost: Extreme Gradient Boosting (gradient-boosted decision-tree algorithm)
Δs: Difference between crop and location silhouette scores (crop-location structure score)
τ² (tau-squared): Between-study (between-protein) heterogeneity variance in meta-analysis
I²: Proportion of total variance attributable to true effect-size heterogeneity
μ (mu): Pooled (random-effects) effect-size estimate
ρ (rho): Spearman rank correlation coefficient
K: Number of classes (crop groups) in multiclass evaluation
SDHI: Succinate dehydrogenase inhibitor (fungicide mode of action)
ROS: Reactive oxygen species
UPS: Ubiquitin-proteasome system
ERAD: Endoplasmic-reticulum-associated degradation
GG-NER: Global-genome nucleotide excision repair
GlcNAc-6P / GlcN-6P: N-acetylglucosamine-6-phosphate / glucosamine-6-phosphate
CCE: Carboxyl/cholinesterase(s)
E1 / E3: Ubiquitin-activating (E1) / ubiquitin-ligase (E3) enzymes
CRL4: Cullin-4 RING E3 ubiquitin ligase
VCP/p97: Valosin-containing protein / p97 (vesicle-fusing ATPase ortholog)
GLUT1: Glucose transporter 1

