## supplementary info for "Comparative Proteomics Across Tissues and Crop Agroecosystems Reveals Agricultural Stressor Responses in the Western Honey Bee"

**Comparative Proteomics Across Tissues and Crop Agroecosystems Reveals Distinct Functional Features**

This PDF file includes:

List of Supplementary Table………………………….………………………….………………………….….….2

Supplementary Figure 1………………………….………………………….………………………….………….3

Supplementary Figure 2………………………….………………………….………………………….………….4

Supplementary Figure 3.………………………….………………………….………………………….…………5

Supplementary Figure 4………………………….………………………….………………………….………….6

Supplementary Figure 5………………………….………………………….………………………….………….7

Supplementary Figure 6………………………….………………………….………………………….……….…8

Supplementary Figure 7………………………….………………………….………………………….………….9

Supplementary Figure 8………………………….………………………….………………………….………...10

Supplementary Figure 9………………………….………………………….………………………….………...11

Supplementary Figure 10………………………….………………………….………………………….……….12

Supplementary Figure 11………………………….………………………….………………………….……….13

Supplementary Figure 12………………………….………………………….………………………….……….15

Supplementary Figure 13………………………….………………………….………………………….……….16

Supplementary Figure 14………………………….………………………….………………………….……….17

Supplementary Figure 15………………………….………………………….………………………….……….18

**List of Supplementary Table (Provided as Additional Excel Files)**

Supplementary Table 1. [Tables with samples and stressor factors](https://docs.google.com/spreadsheets/d/12OwC1uq46khV4PEM02LUKEghPU5Czn5J6E4wLy-vCHo/edit?gid=281628786#gid=281628786)

Supplementary Table 2. List of abdomen-specific protein names and their accessions

Supplementary Table 3. List of gut-specific protein names and their accessions

Supplementary Table 4. List of head-specific protein names and their accessions

Supplementary Table 5. Random Effect Meta-analysis for crop-dominant effects across tissues and years

Supplementary Table 6. Environmental burden profiles (pesticides) per crop and year

Supplementary Table 7. Environmental burden profiles (virus) per crop and year

Supplementary Table 8. Environmentally sensitive proteins

Supplementary Table 9. Significant stressors (per-term ANOVA)

Supplementary Table 10. Spearman correlations passing FDR < 0.05 in 2020.

Supplementary Table 11. Proteins of different origins showing reproducible negative association with Boscalid across years in the gut or head

a)
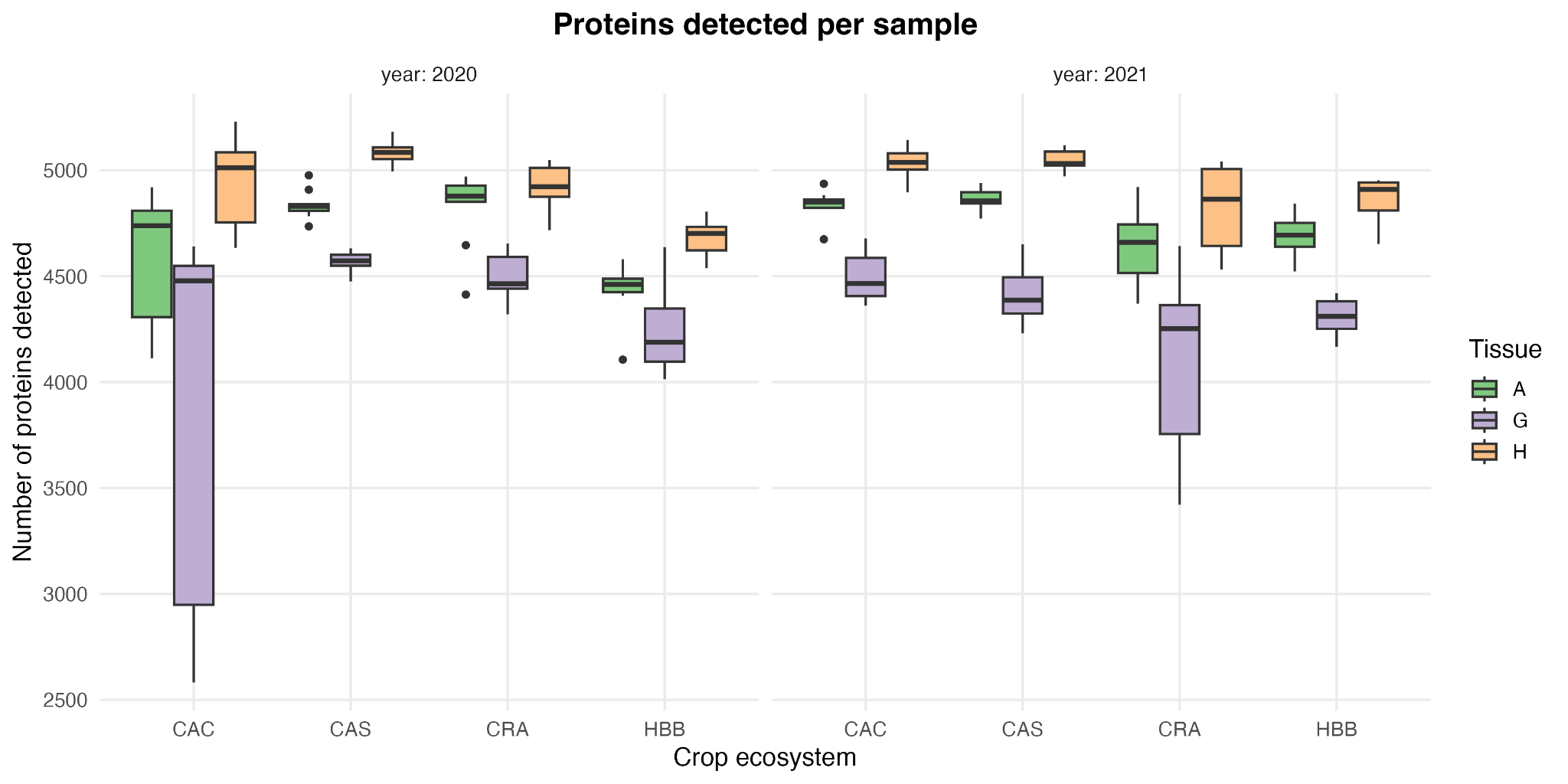

b)
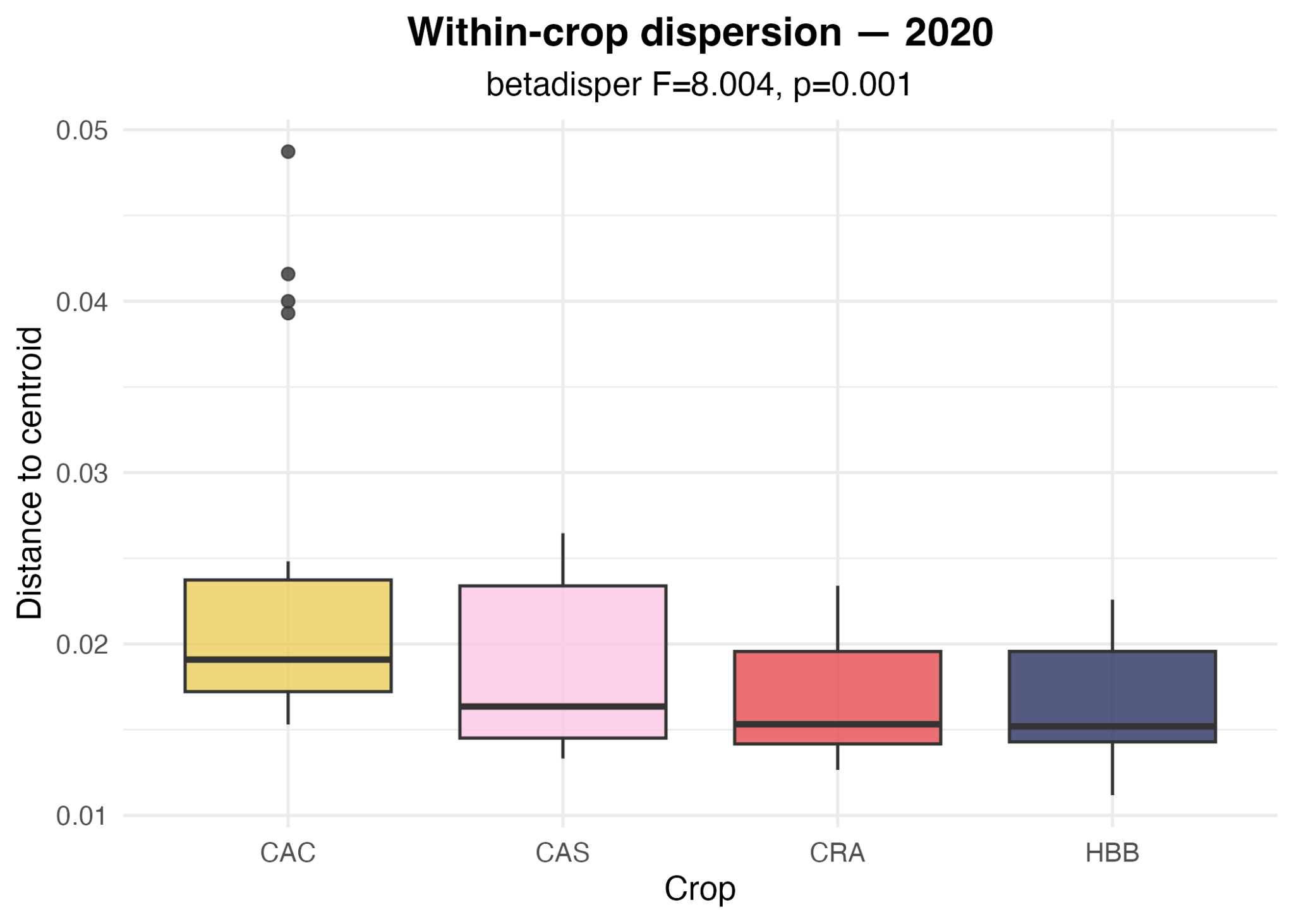

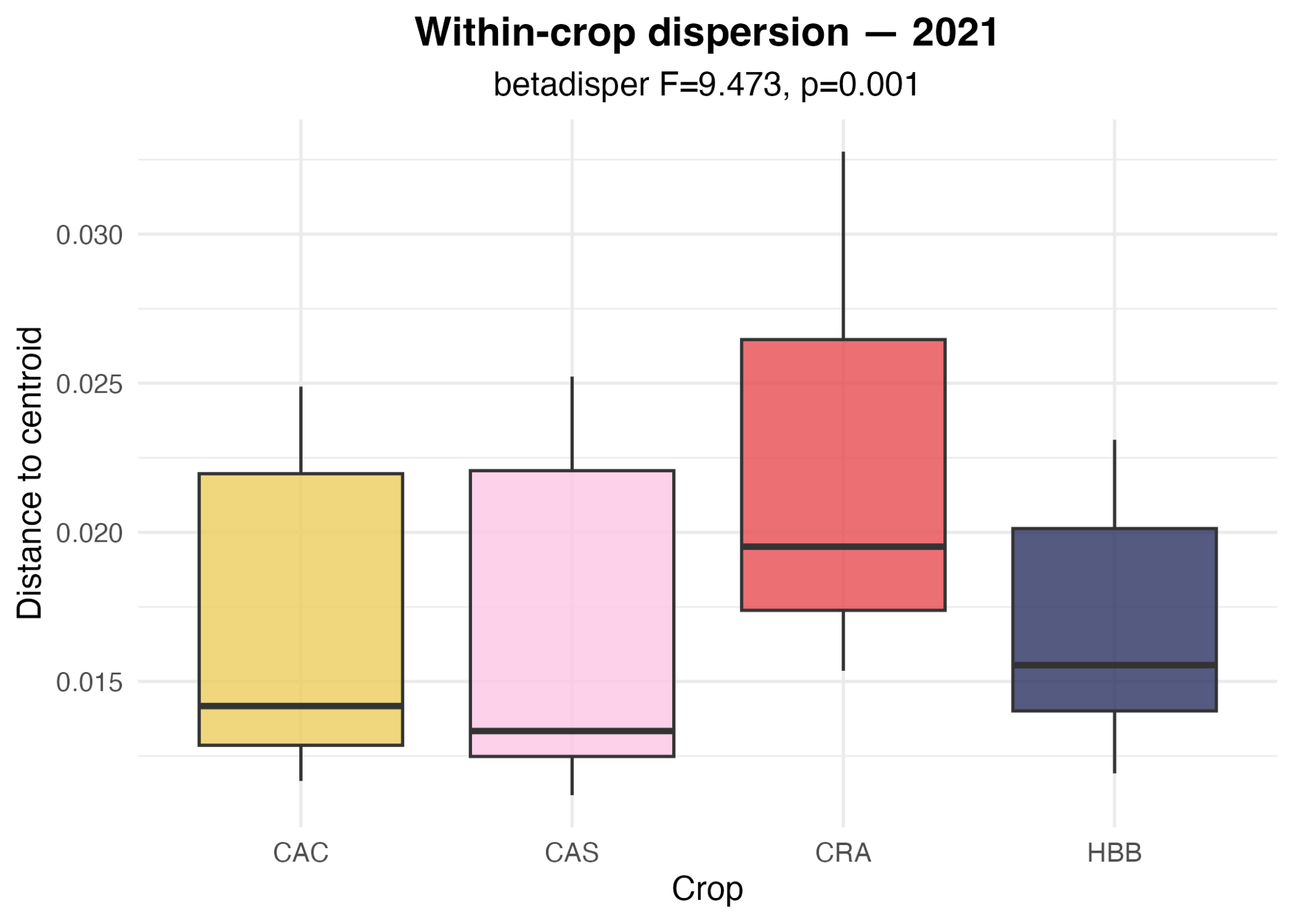

c)
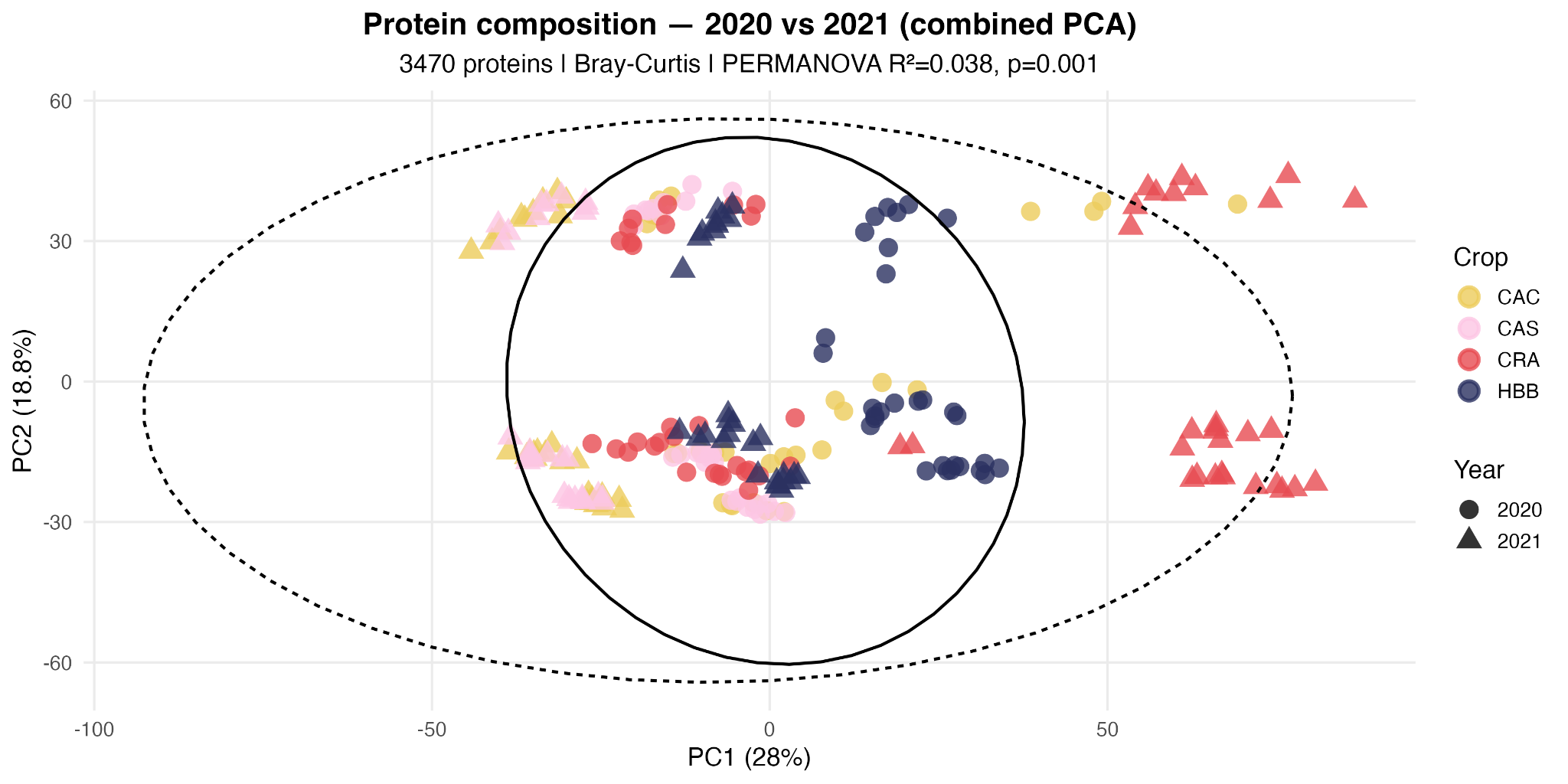

d)
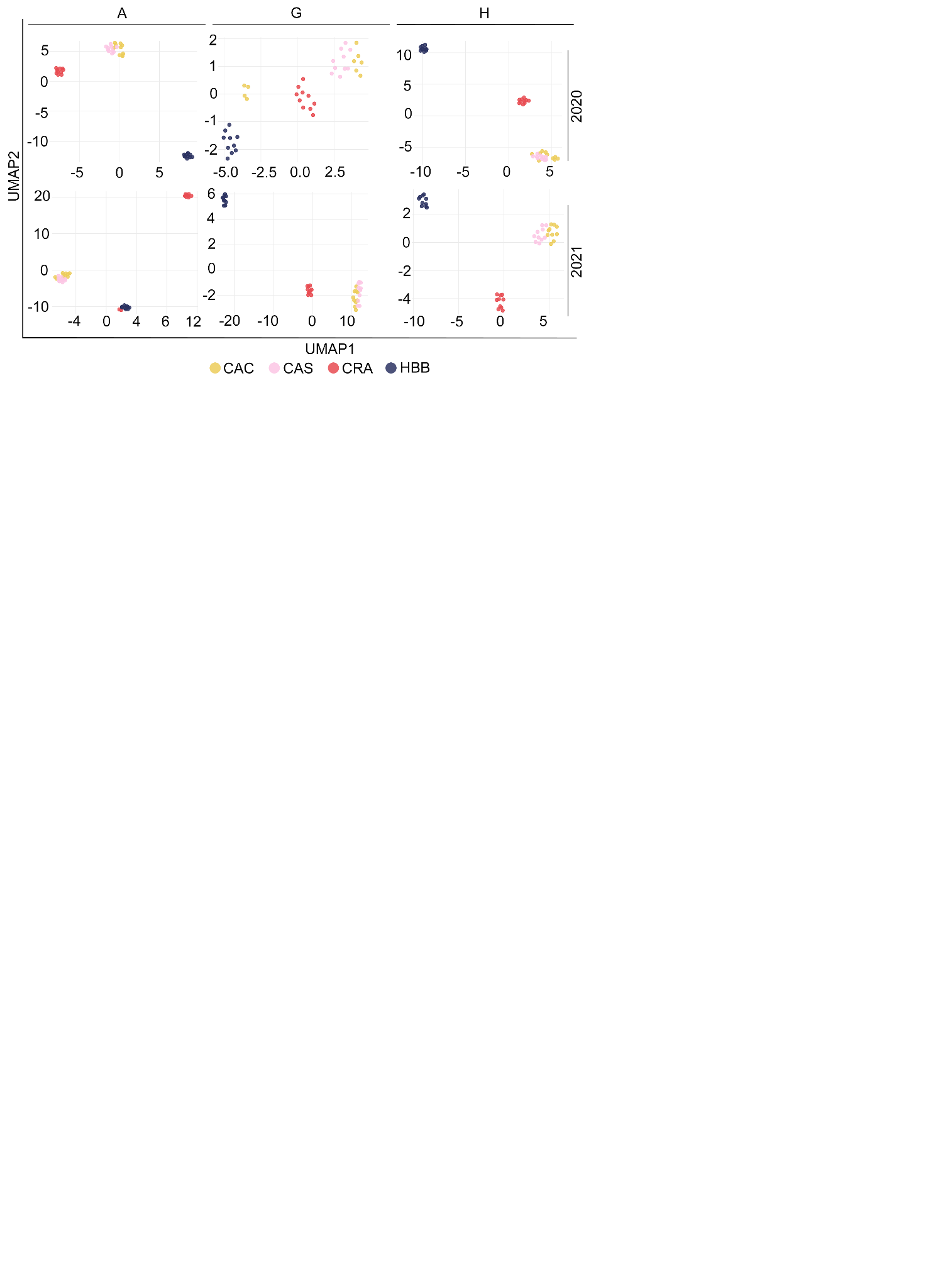

**Supplementary Figure 1. Crop structure and dispersion in proteomic space**

**(a)** Distribution of the number of detected proteins per sample across crop ecosystems, tissues, and years. Boxplots summarize proteome coverage for each crop-tissue combination and were used to assess sample-level missingness and potential systematic differences in protein detection depth across experimental groups. **(b)** Betadisper analysis showing within-crop dispersion differences across years. Dispersion heterogeneity is significant in both 2020 (F = 8.00, p = 0.001) and 2021 (F = 9.47, p = 0.001) **(c)** UMAP. **(d)** PCA of Bray-Curtis distances for 2020 (PC1 = 21.6%, PC2 = 17.8%) and 2021 (PC1 = 40.6%, PC2 = 19.5%). Crop identity explains significant variation in both years (PERMANOVA R^2^ = 0.226 and 0.498, p = 0.001). Tissue is indicated by shape.

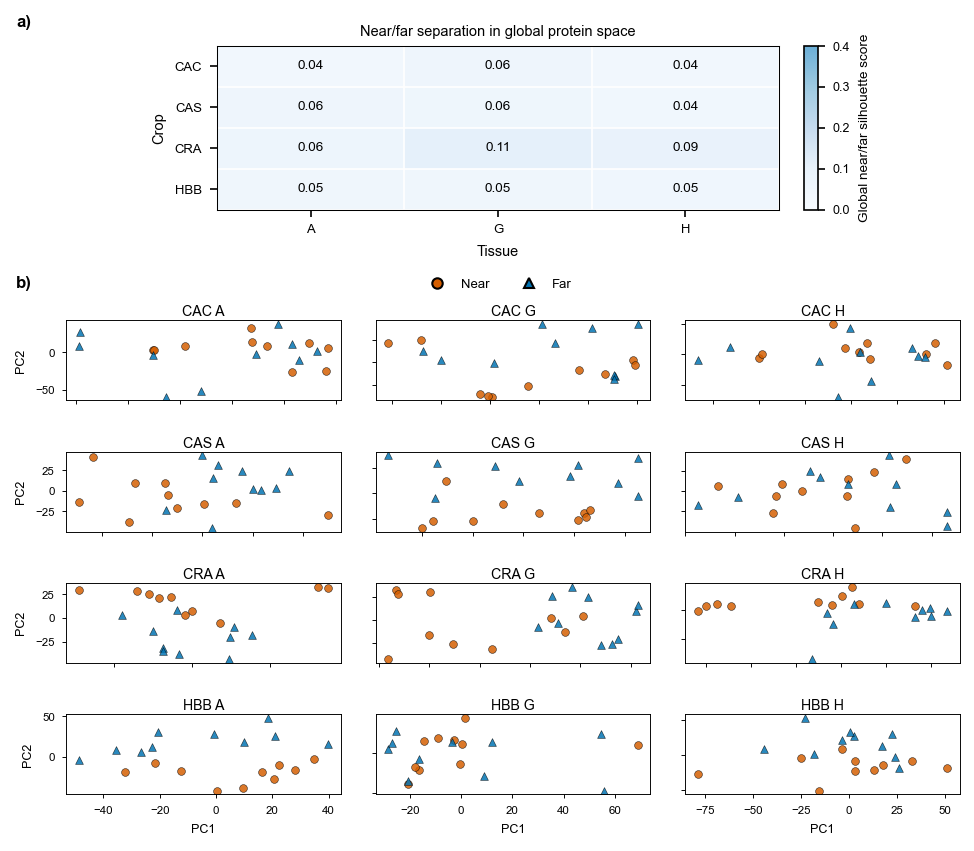

**Supplementary Figure 2. Near/far exposure sensitivity analysis in the global protein expression space.**

**(a)** Heatmap of near/far silhouette scores computed within each crop-by-tissue subset after year-wise adjustment. **(b)** PCA ordinations for each crop-by-tissue subset, with samples coloured and shaped by near/far exposure status. Consistently low silhouette scores and substantial overlap between near and far samples indicate that near/far placement contributed only weakly to global proteomic variation and did not define a dominant separable subgroup.

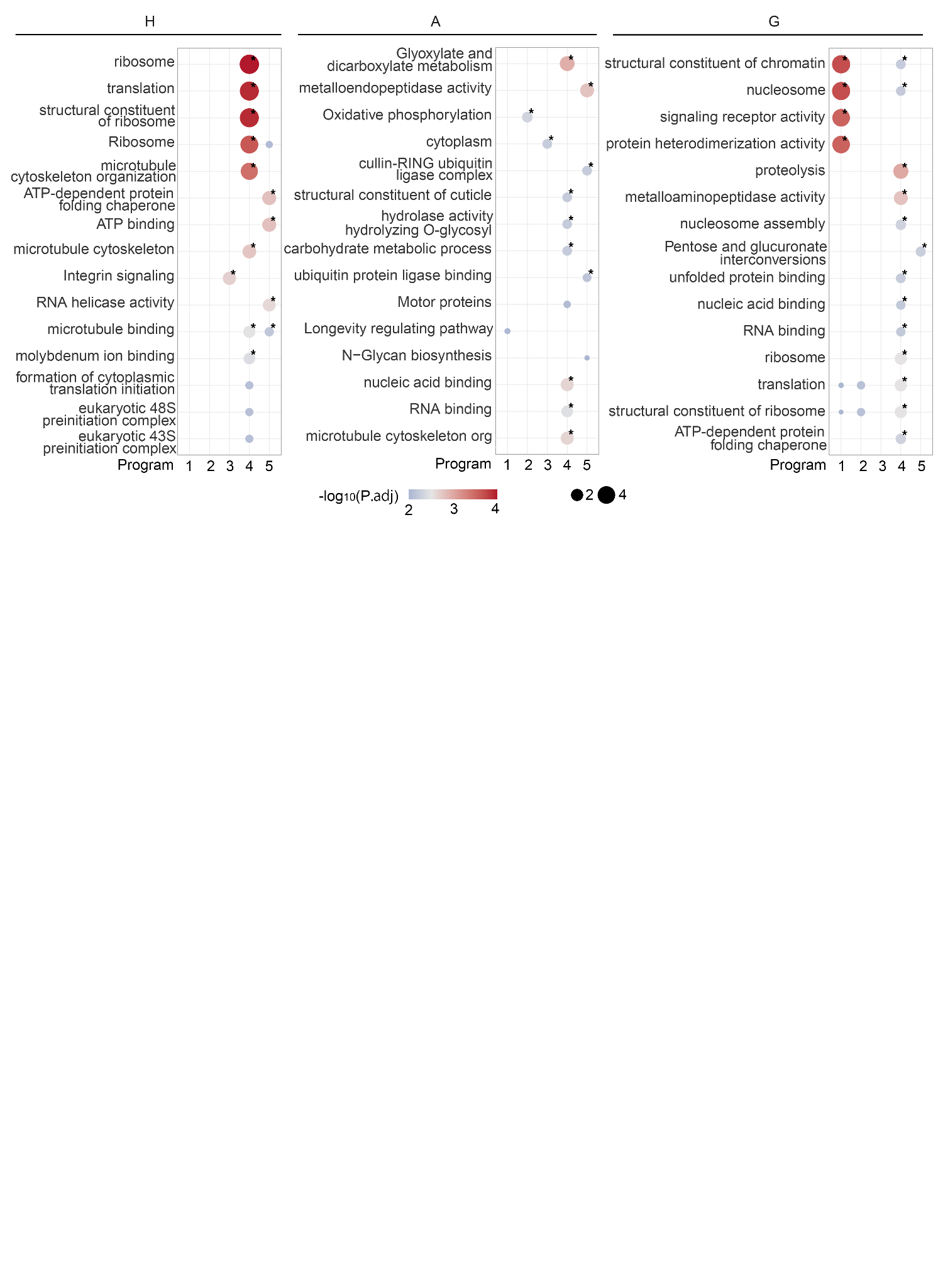
**Supplementary Figure 3.** Gene Ontology (GO) enrichment analysis of proteins with high loadings in tissue-specific latent programs. Dot size indicates enrichment significance (-log_10_(P)), and colour represents enrichment score.

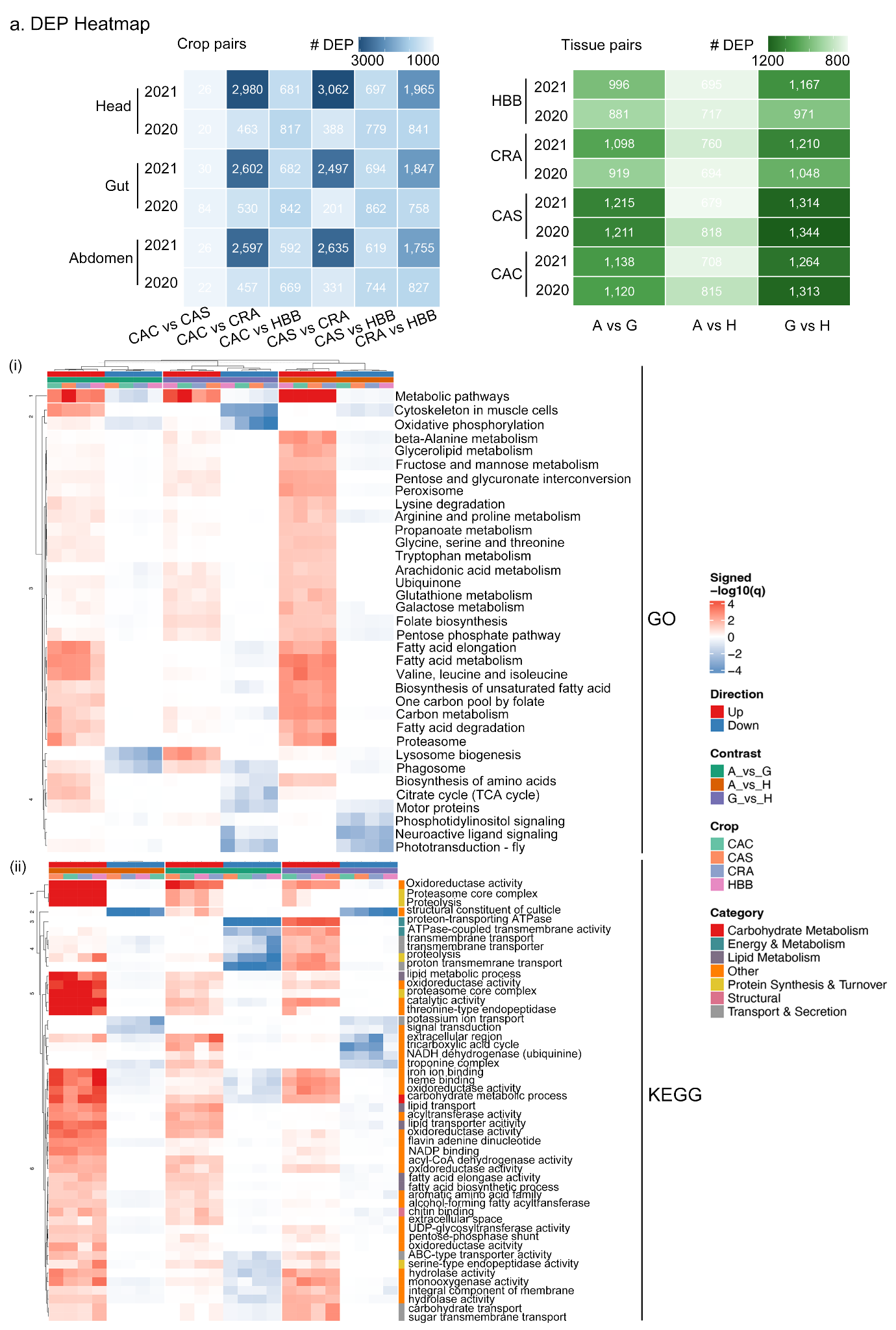
**Supplementary Figure 4.** Heatmap showing (a) GO and (b) KEGG enrichment patterns of tissue-specific DEPs across tissue pairwise contrasts. Color intensity represents scaled enrichment scores/logFC values, where red indicates relative enrichment/upregulation and blue indicates relative depletion/downregulation.

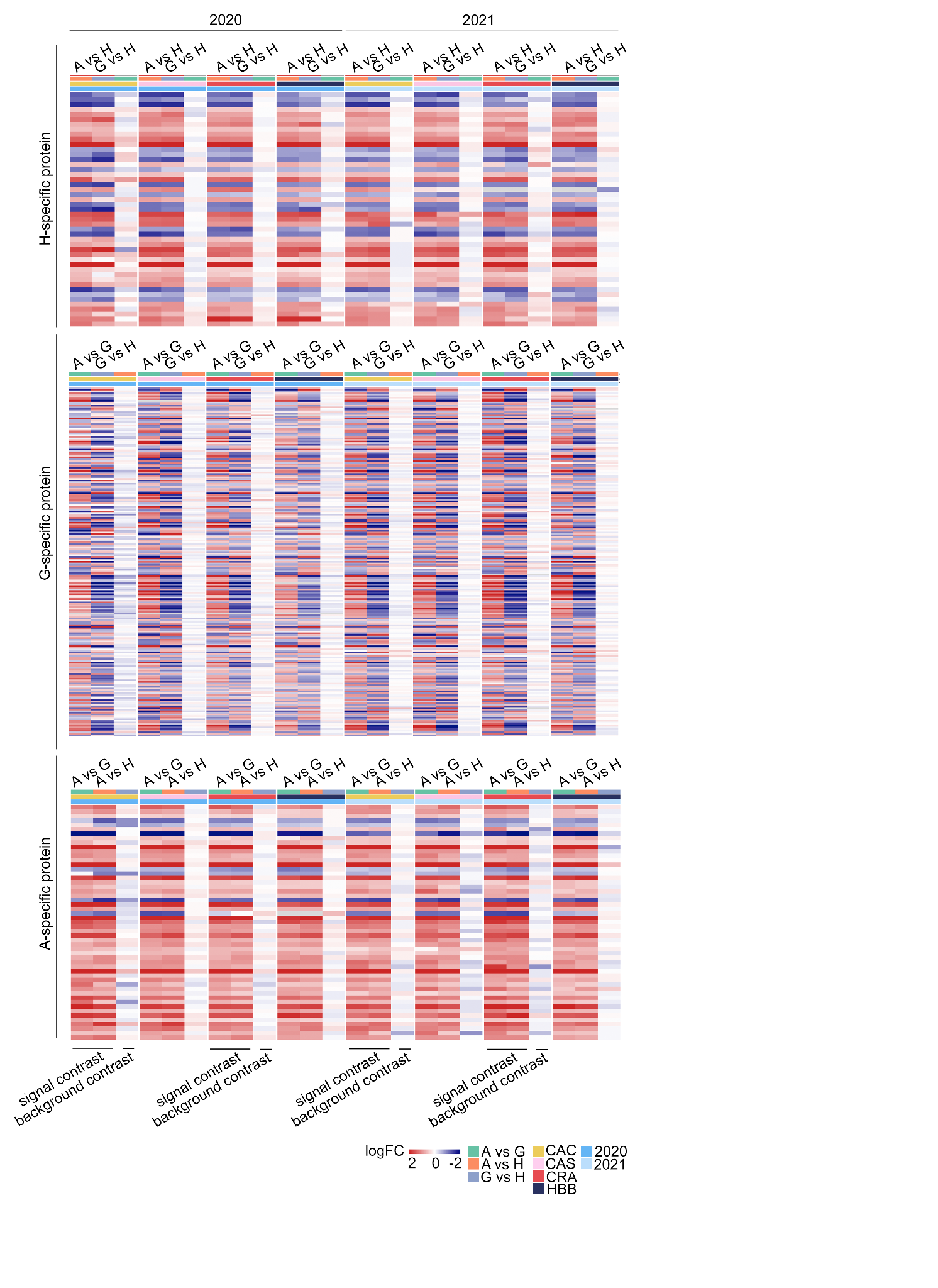

**Supplementary Figure 5.** Heatmap showing scaled expression patterns (logFC) of tissue-specific proteins across tissue contrasts (A vs G, A vs H, and G vs H) in different crops and years. Red and blue indicate positive and negative logFC values, respectively.

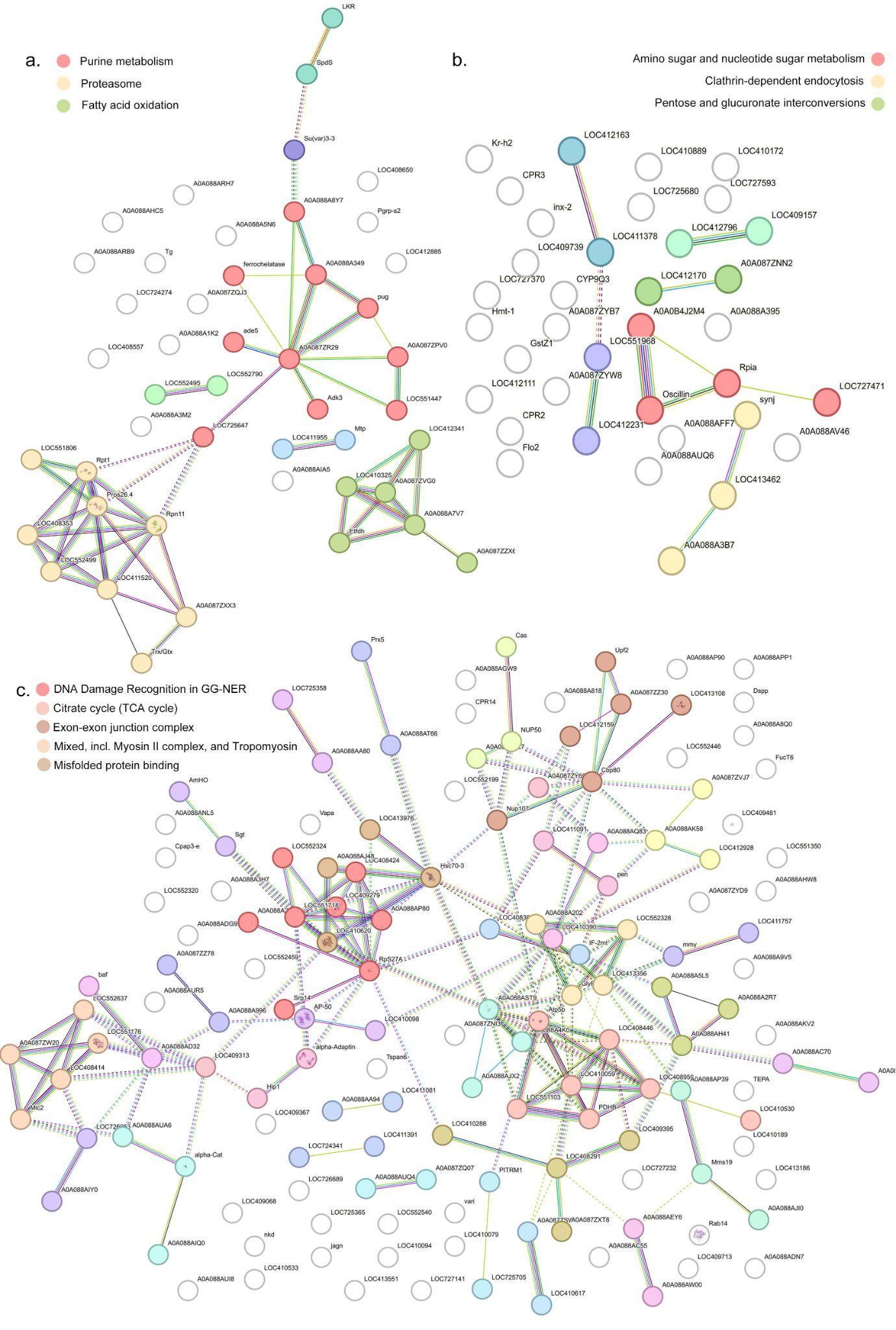

**Supplementary Figure 6.** Global PPI networks constructed from tissue-specific DEPs using STRING analysis. Nodes represent proteins and edges indicate predicted or experimentally supported protein-protein associations.

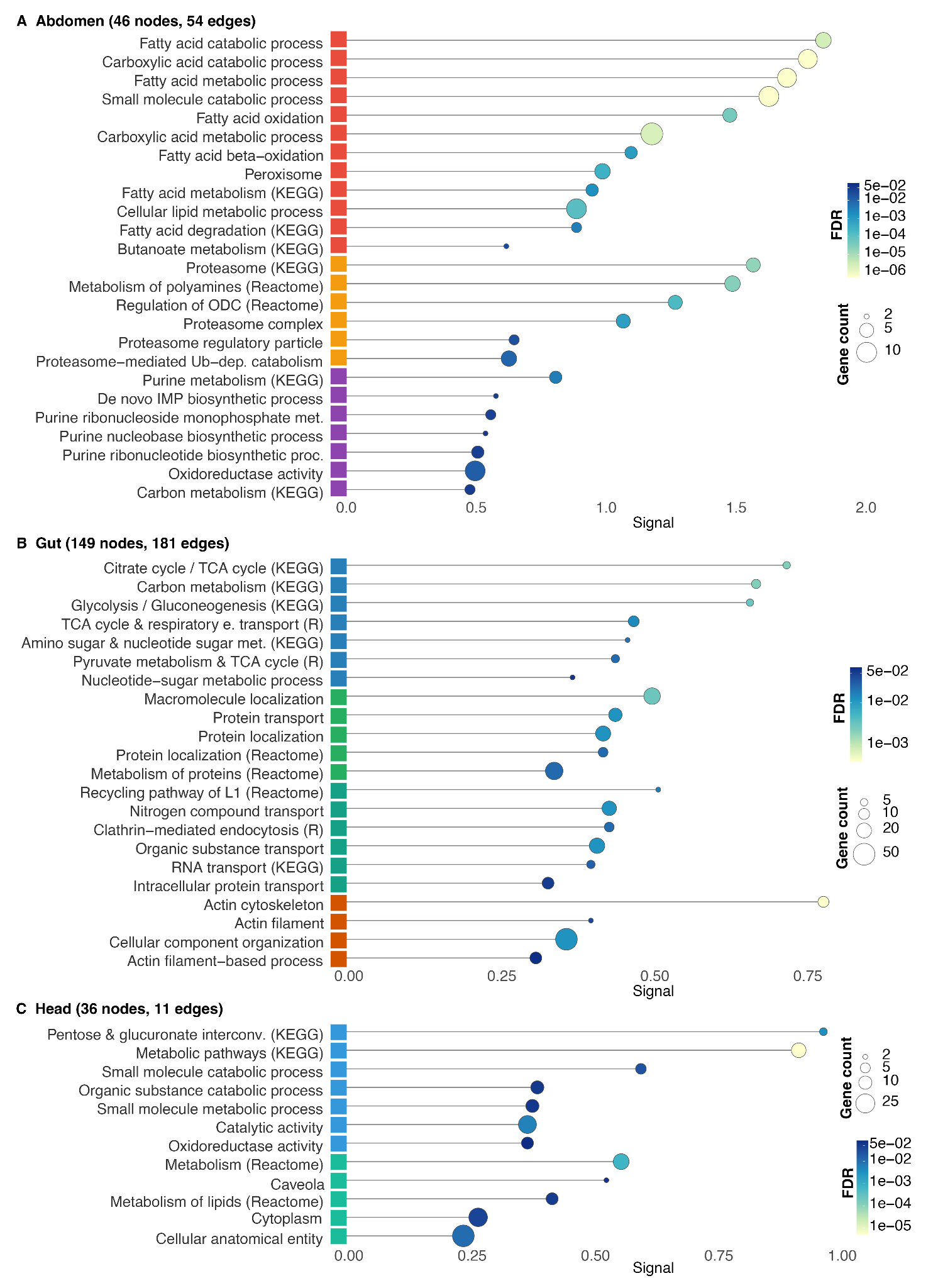

**Supplementary Figure 7.** Functional enrichment bubble plots of tissue-specific PPI subnetworks in abdomen, gut, and head tissues. Bubble size represents the number of proteins associated with each enriched term, and bubble color indicates false discovery rate (FDR)-adjusted significance. Only significantly enriched terms (FDR < 0.05) are shown.

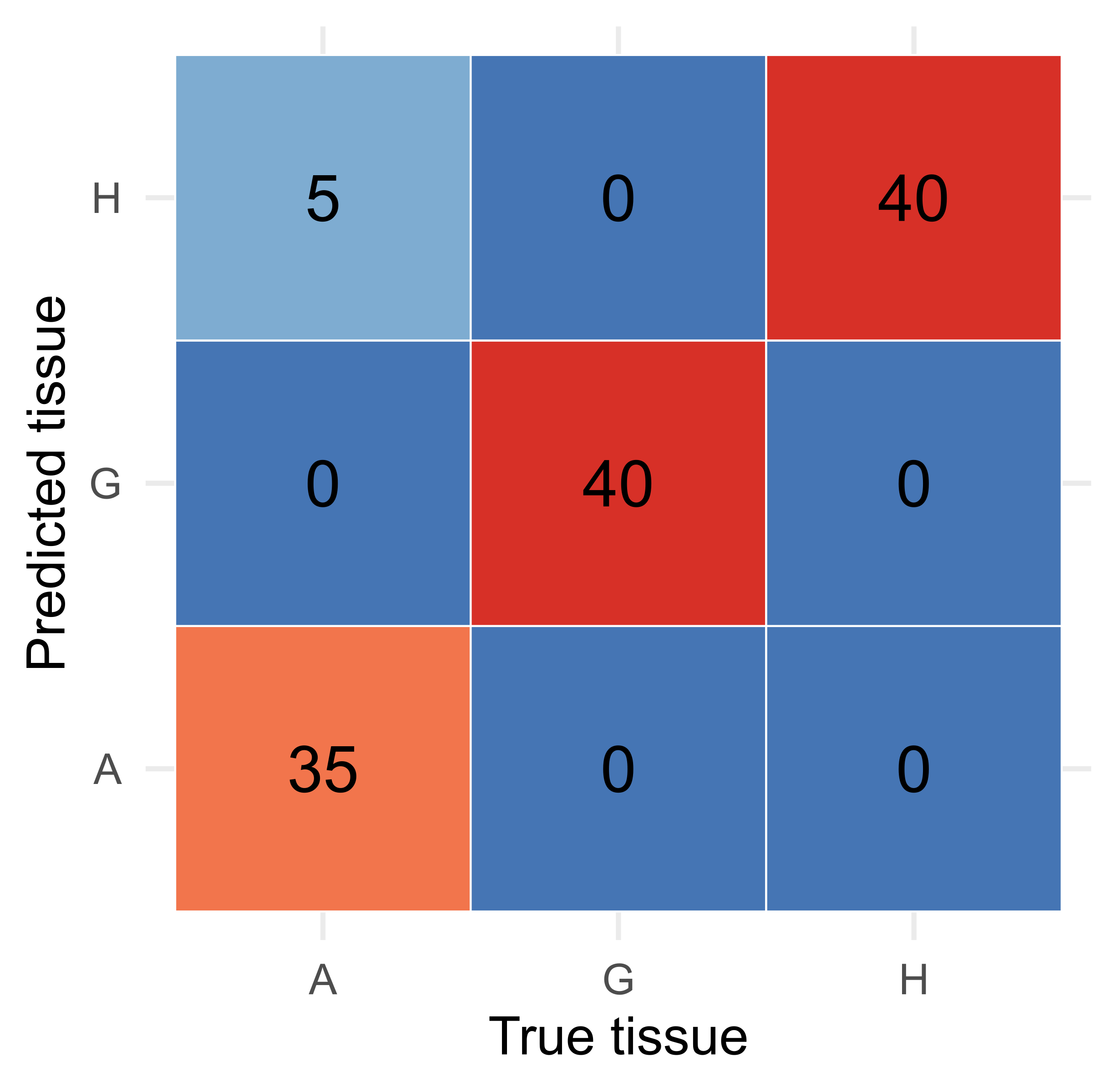

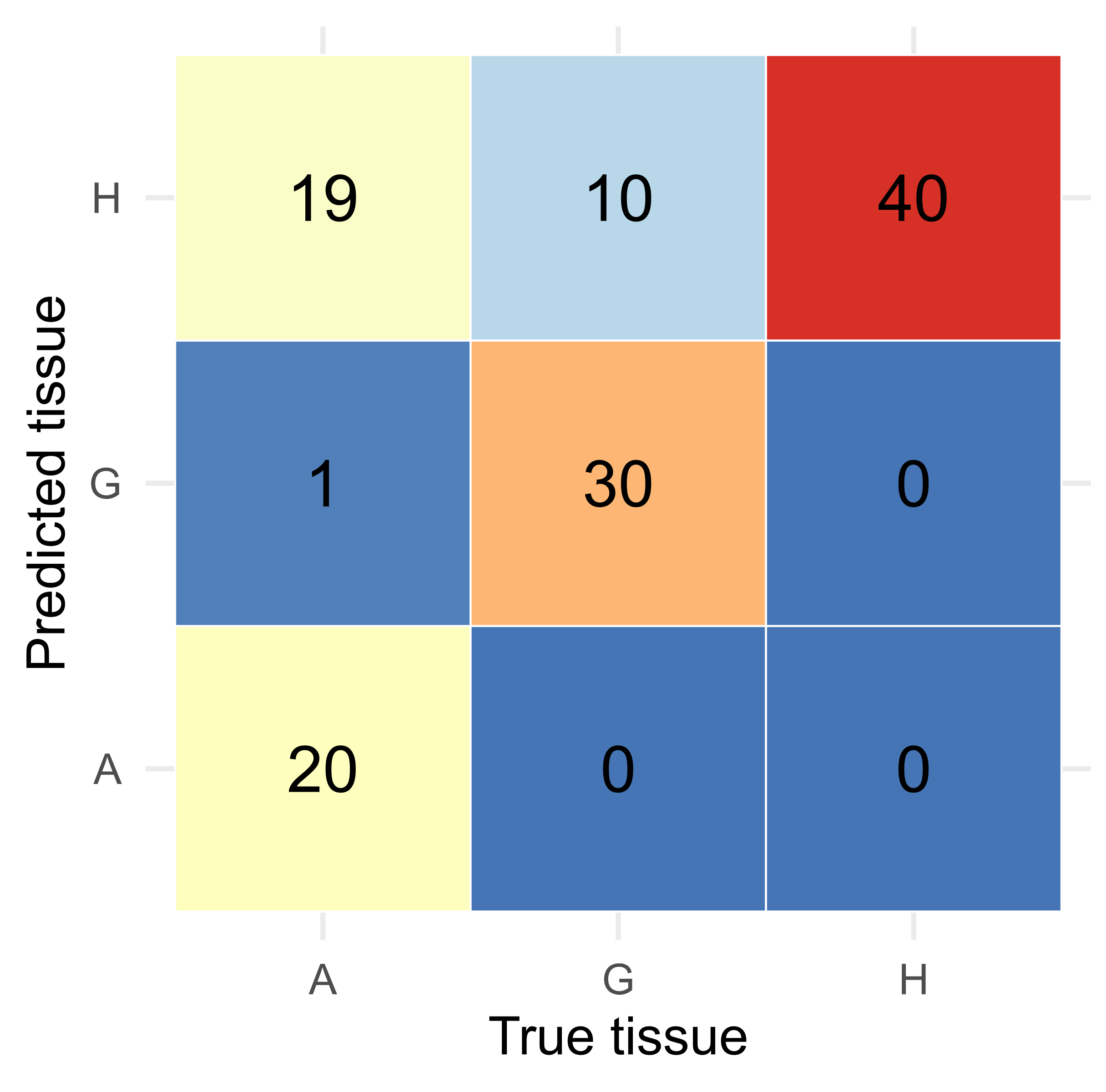

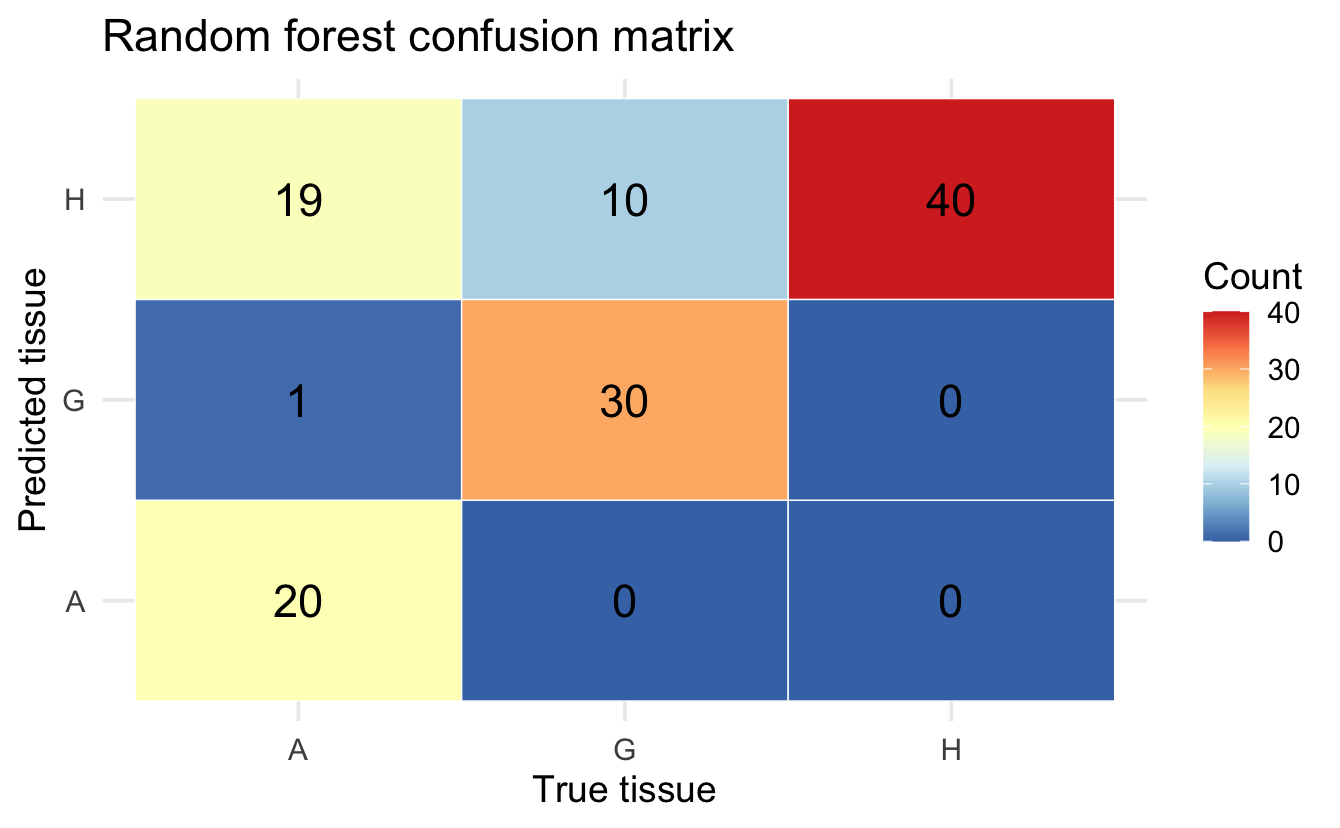

**i	ii**

**Supplementary Figure 8.** Confusion matrices showing bee tissue predictions for the 2021 dataset using a random forest **(i)** and logistic regression **(ii)** models trained on the 2020 dataset. The model was built using limma-selected features shared between both years.

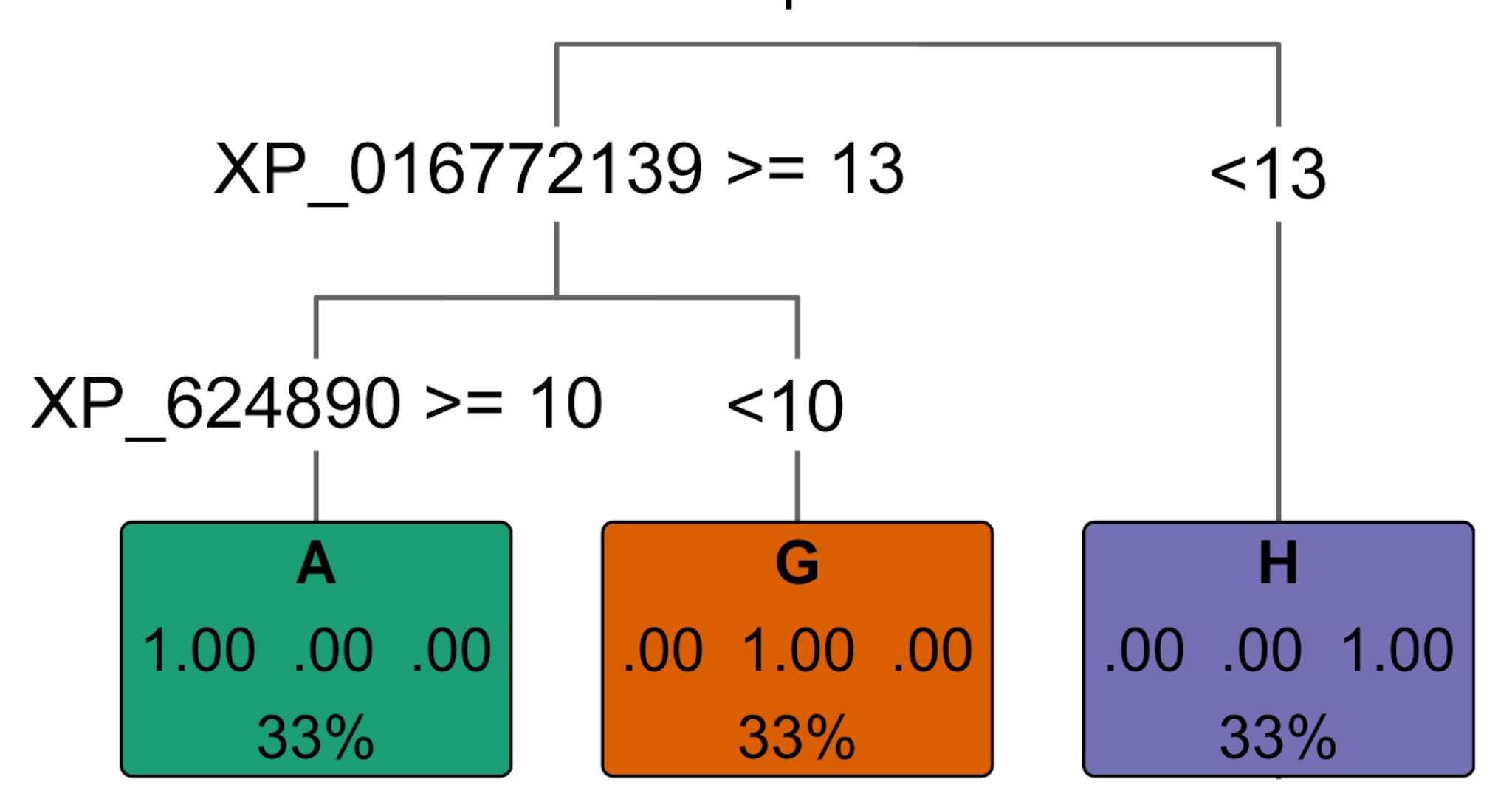

**Supplementary Figure 9.** Decision tree, built based on top random forest features

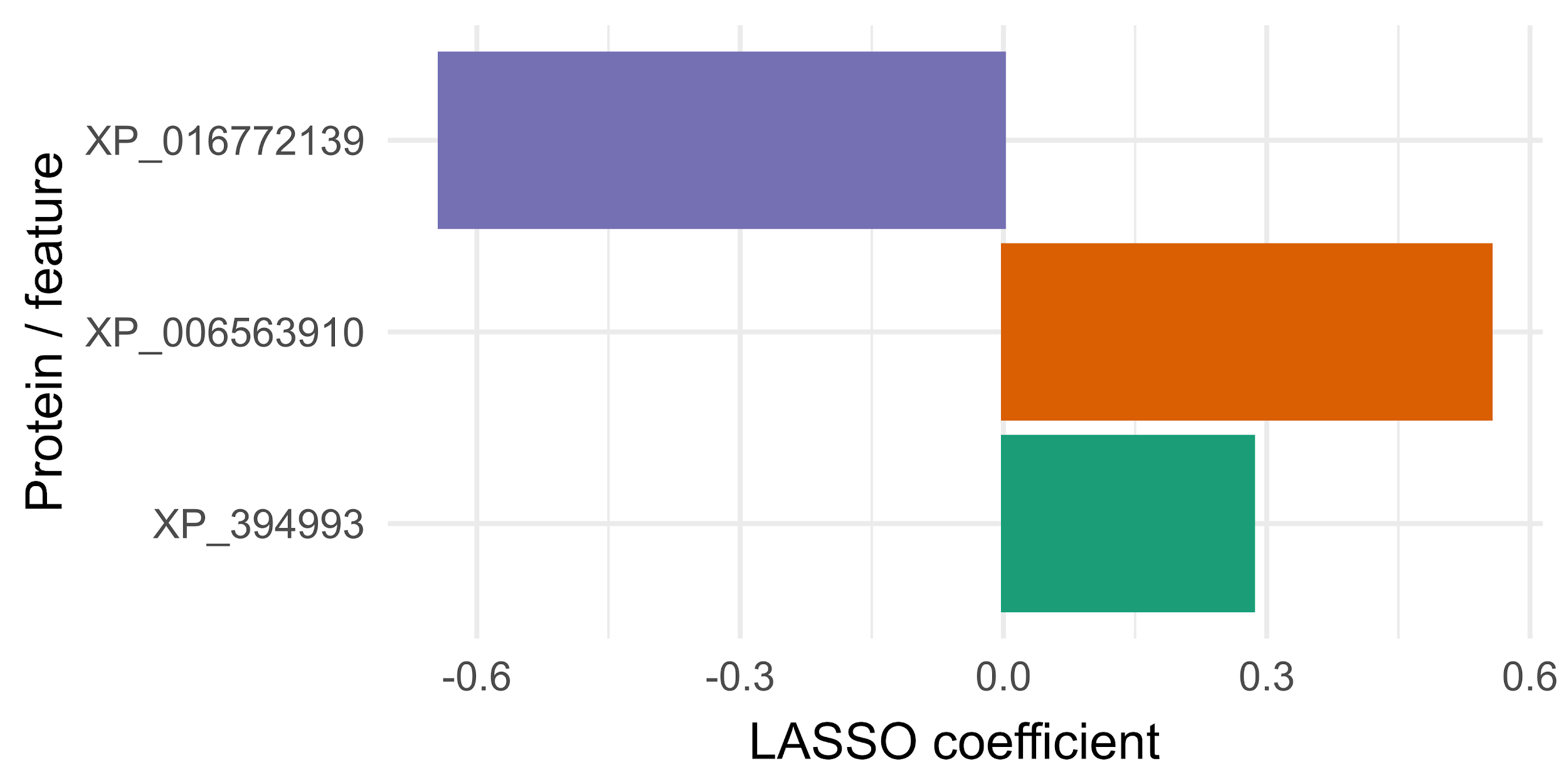

**Tissue**

**H**

**G**

**A**

**Supplementary Figure 10.** Bar plot shows proteins selected by multinomial LASSO logistic regression for bee tissue classification. Bar height and direction represent the LASSO coefficient for each selected protein, with colors indicating the associated tissue class.

(a)
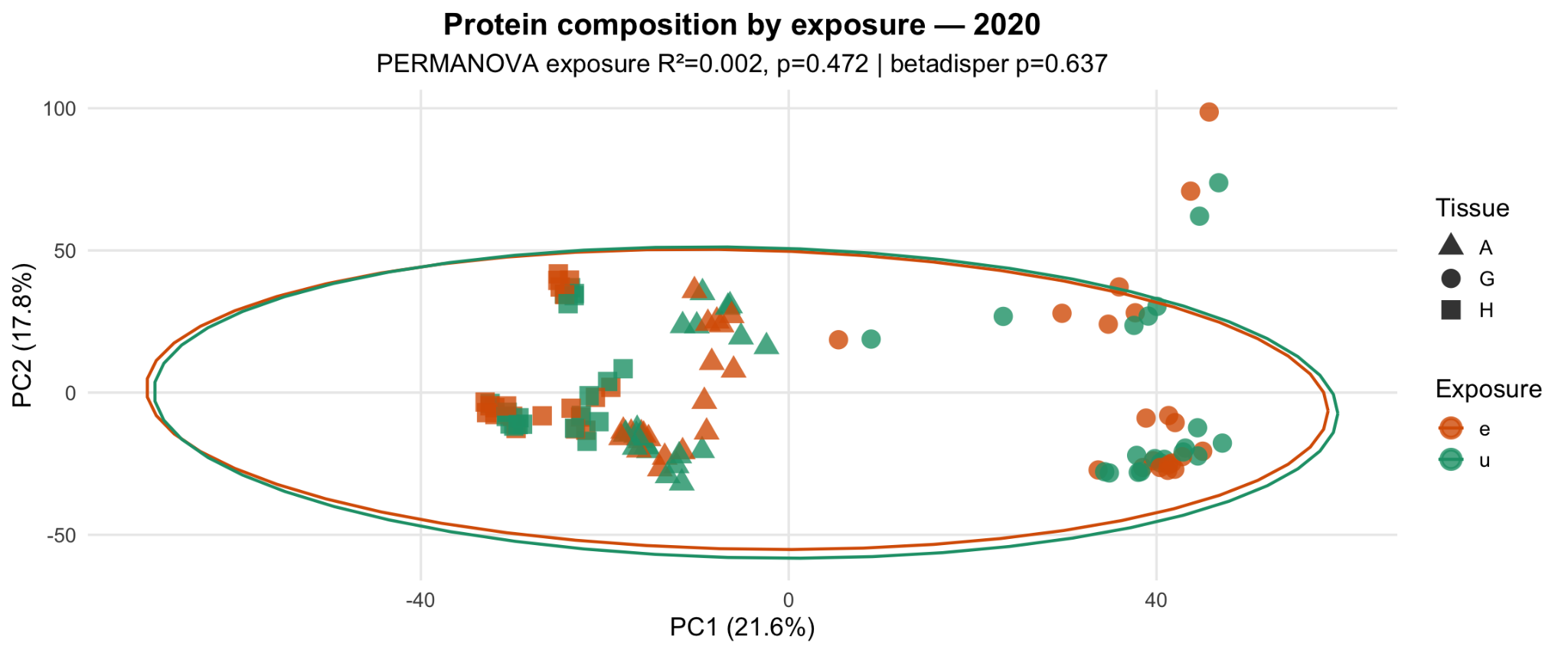

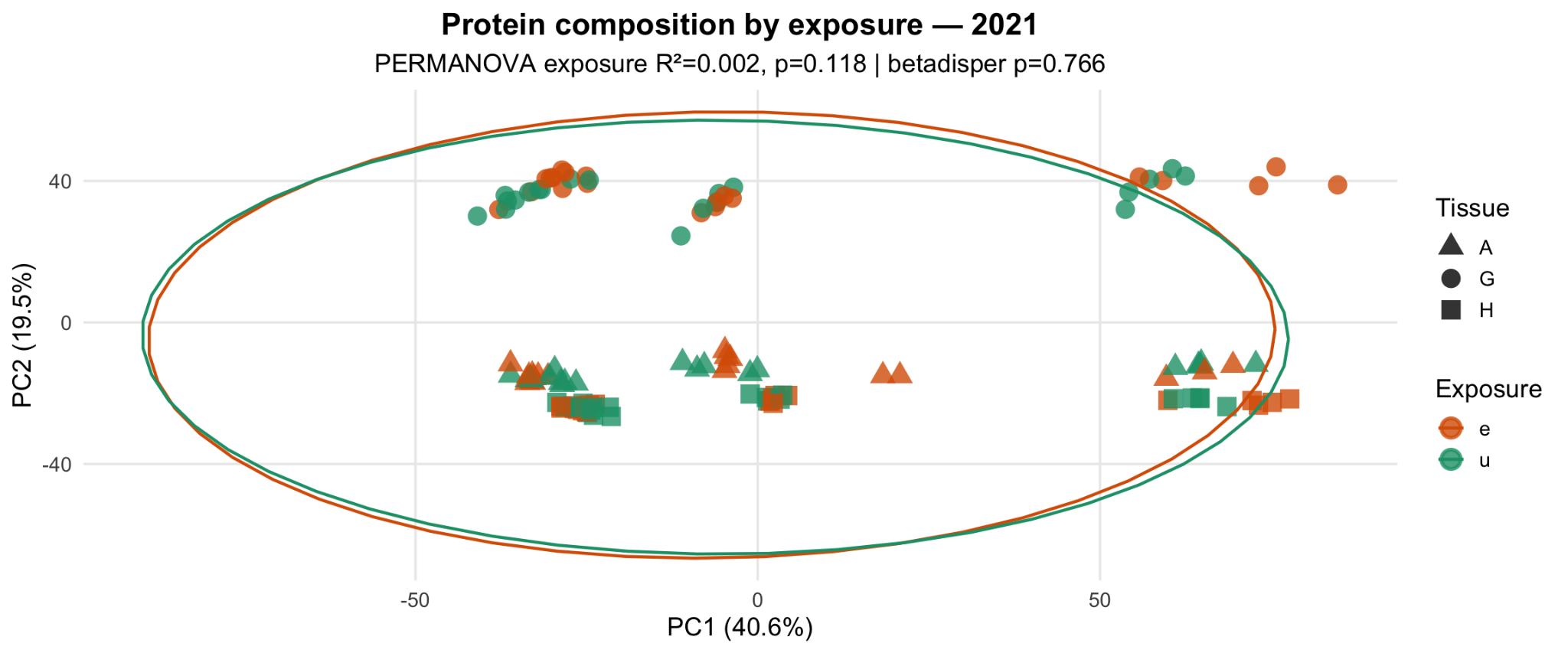

(b)
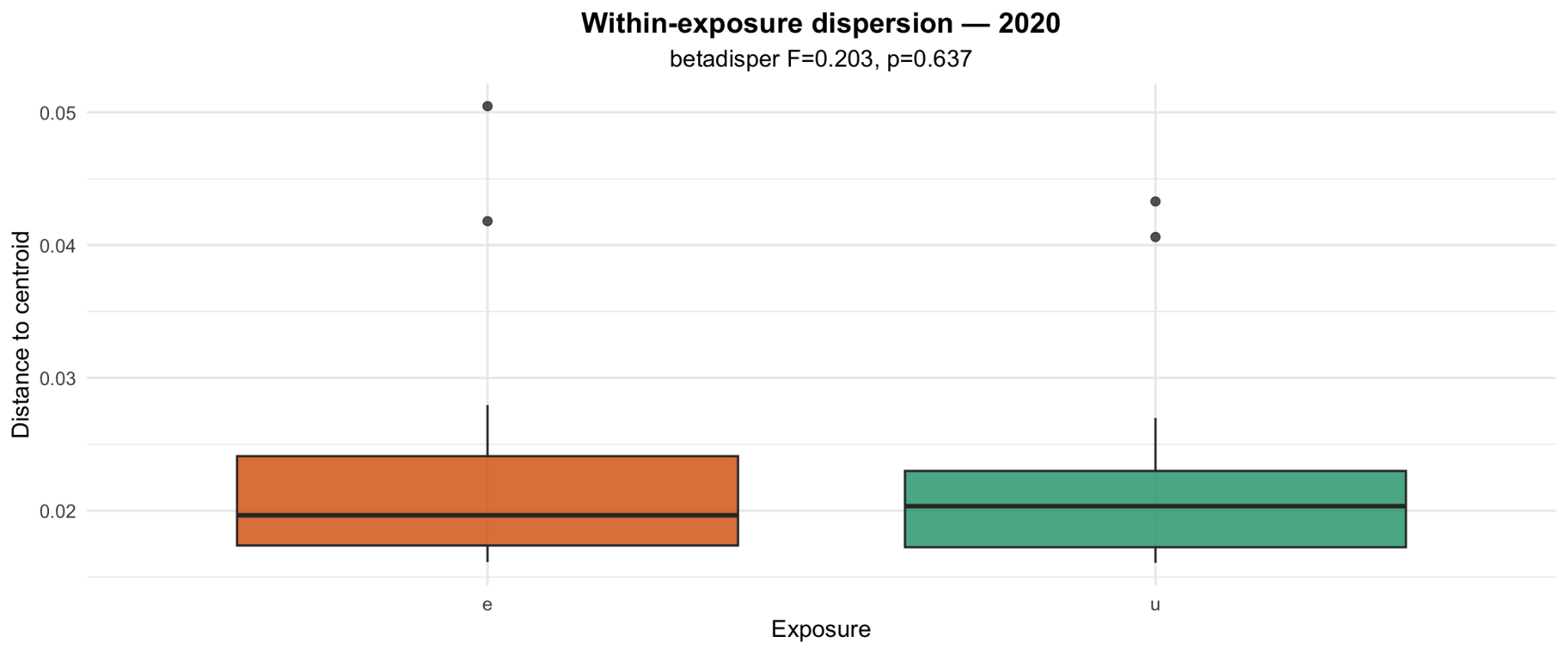

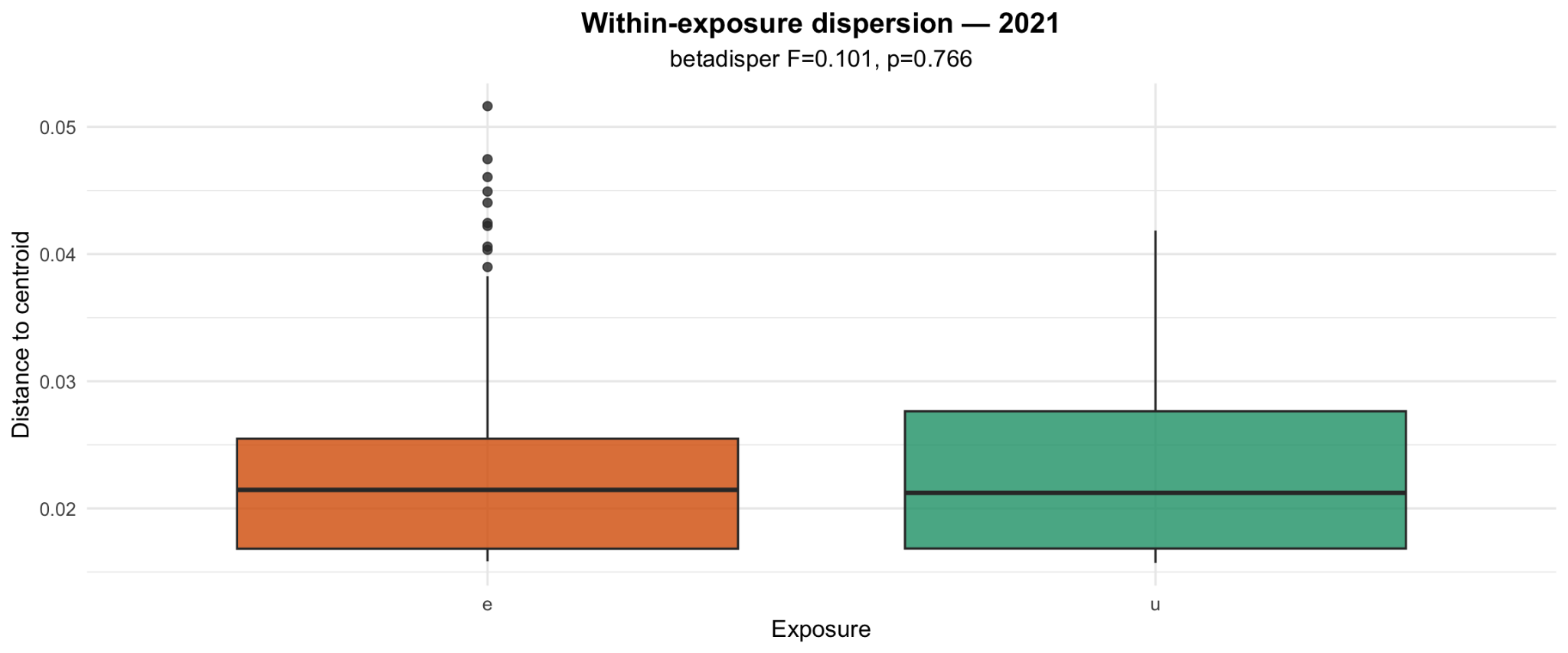

**Supplementary Figure 11.** PERMANOVA **(a)** and betadisper **(b)** results for pesticide exposure effect on protein composition

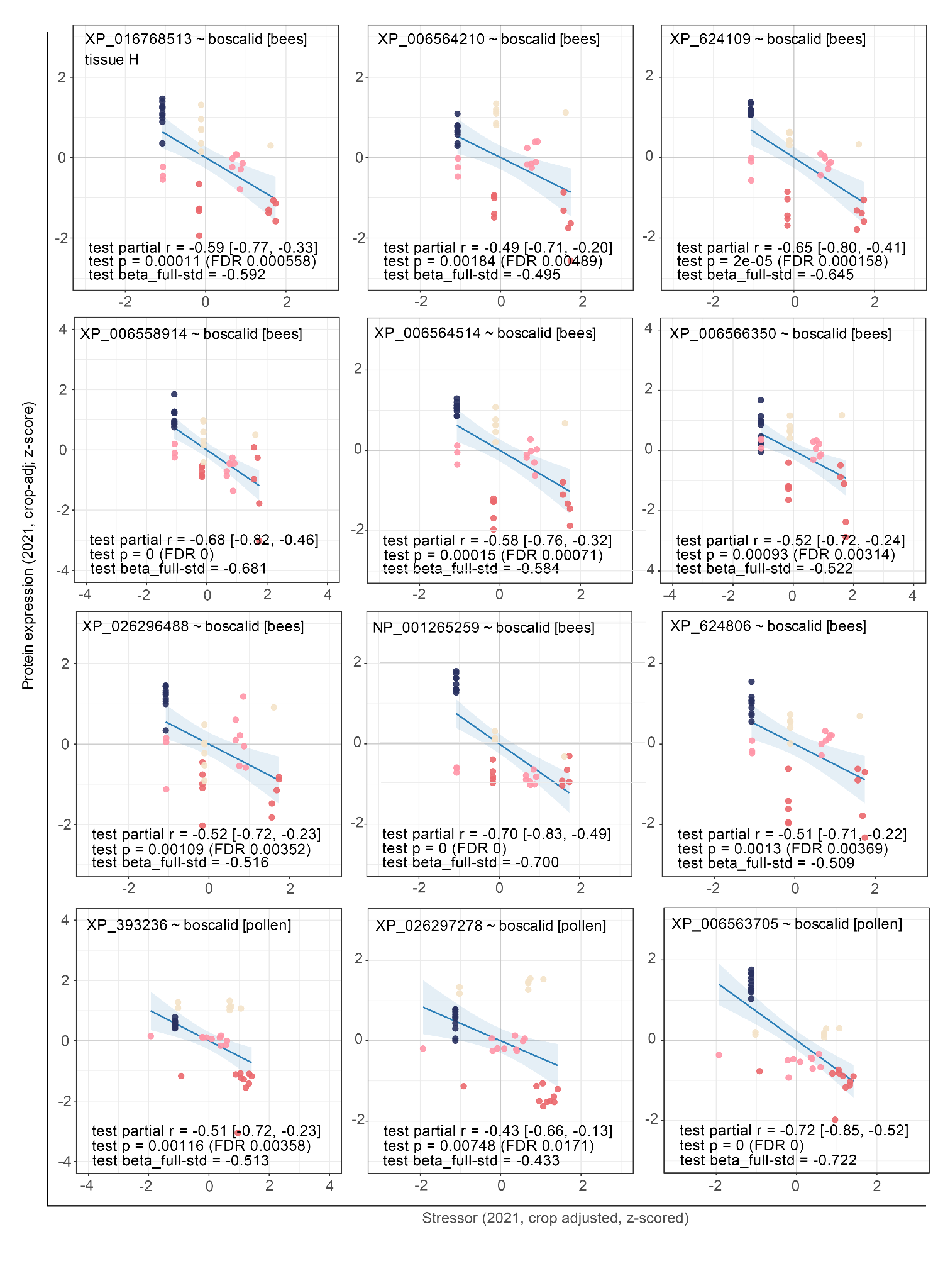

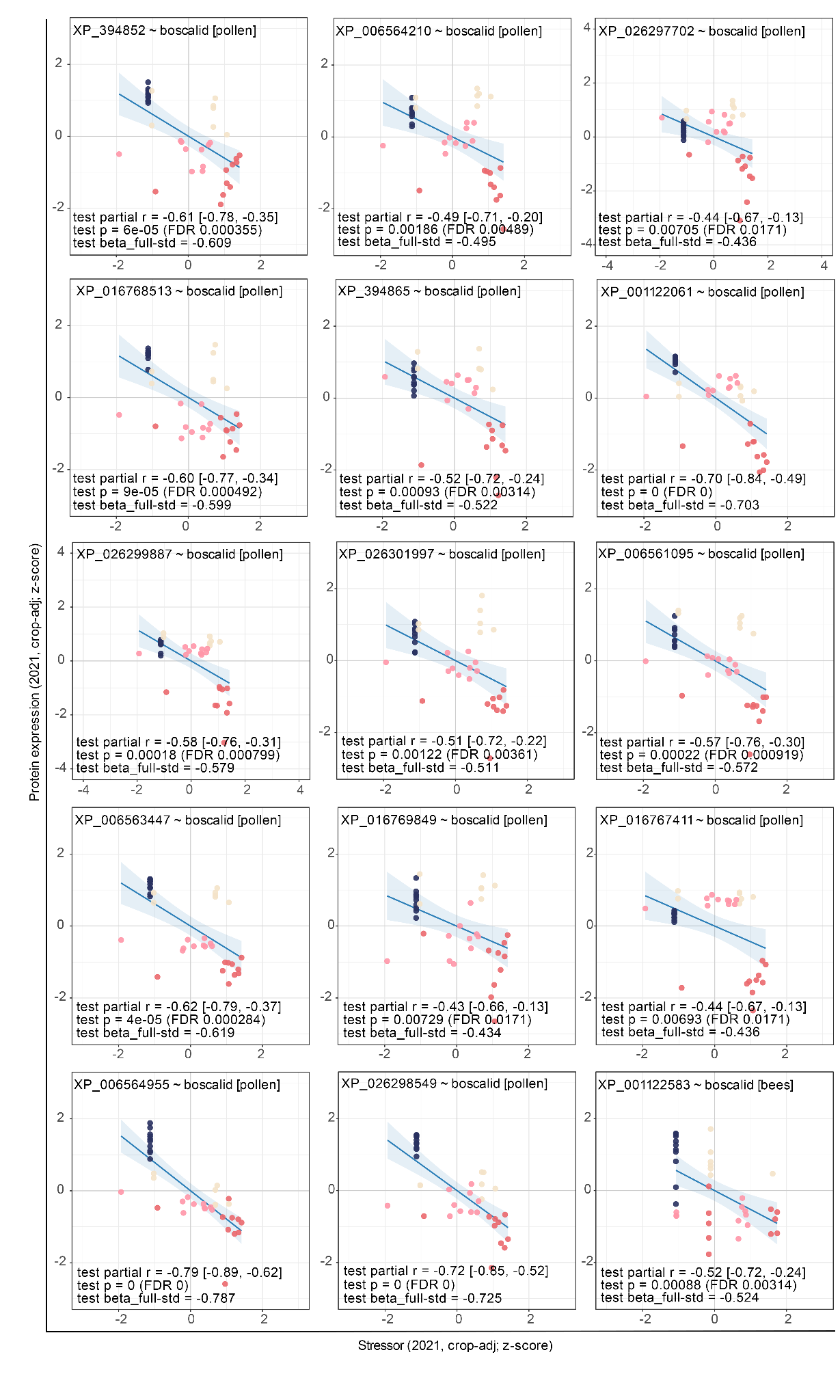
**Supplementary Figure 12.** Representative 2020 protein-stressor association (selected after envfit pre-filtering) independently reproduced in 2021 using sign-consistent FDR-significant validation criteria. There are 26 gut proteins (unspecified in the graph) and 1 head protein (specified in the graph).

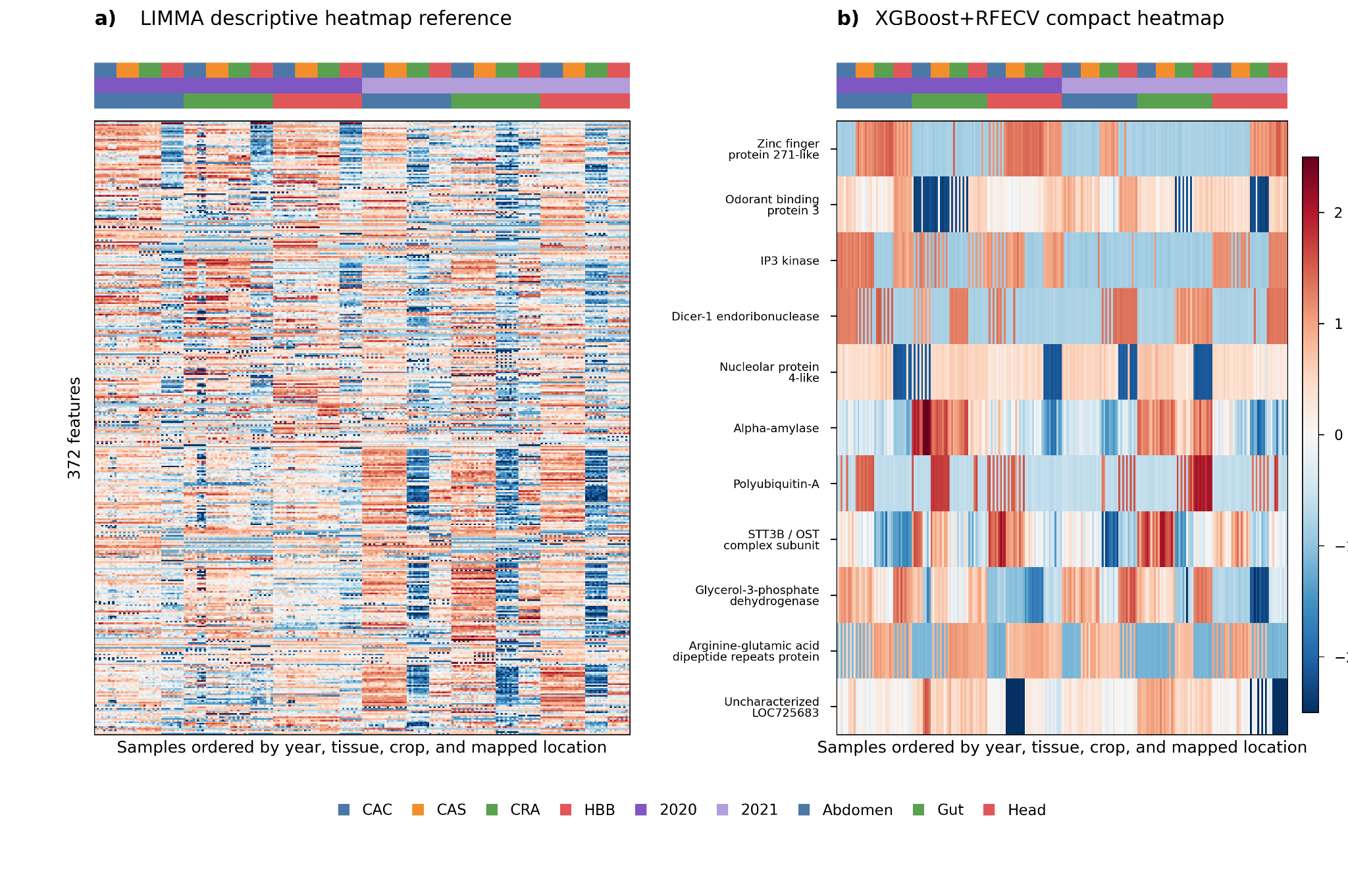

**Supplementary Figure 13. Heatmap comparing broad LIMMA-derived descriptive features with compact XGBoost+RFECV-selected protein features.**

**(a)** LIMMA descriptive reference heatmap containing 372 unique proteins aggregated across tissue-year crop contrasts after duplicate removal. **(b)** XGBoost+RFECV compact heatmap containing 11 selected proteins across abdomen, gut, and head. Columns represent samples ordered by year, tissue, crop, and mapped location; rows represent proteins. This comparison illustrates the shift from broad univariate descriptive structure to sparse supervised feature panels used for temporal prediction.

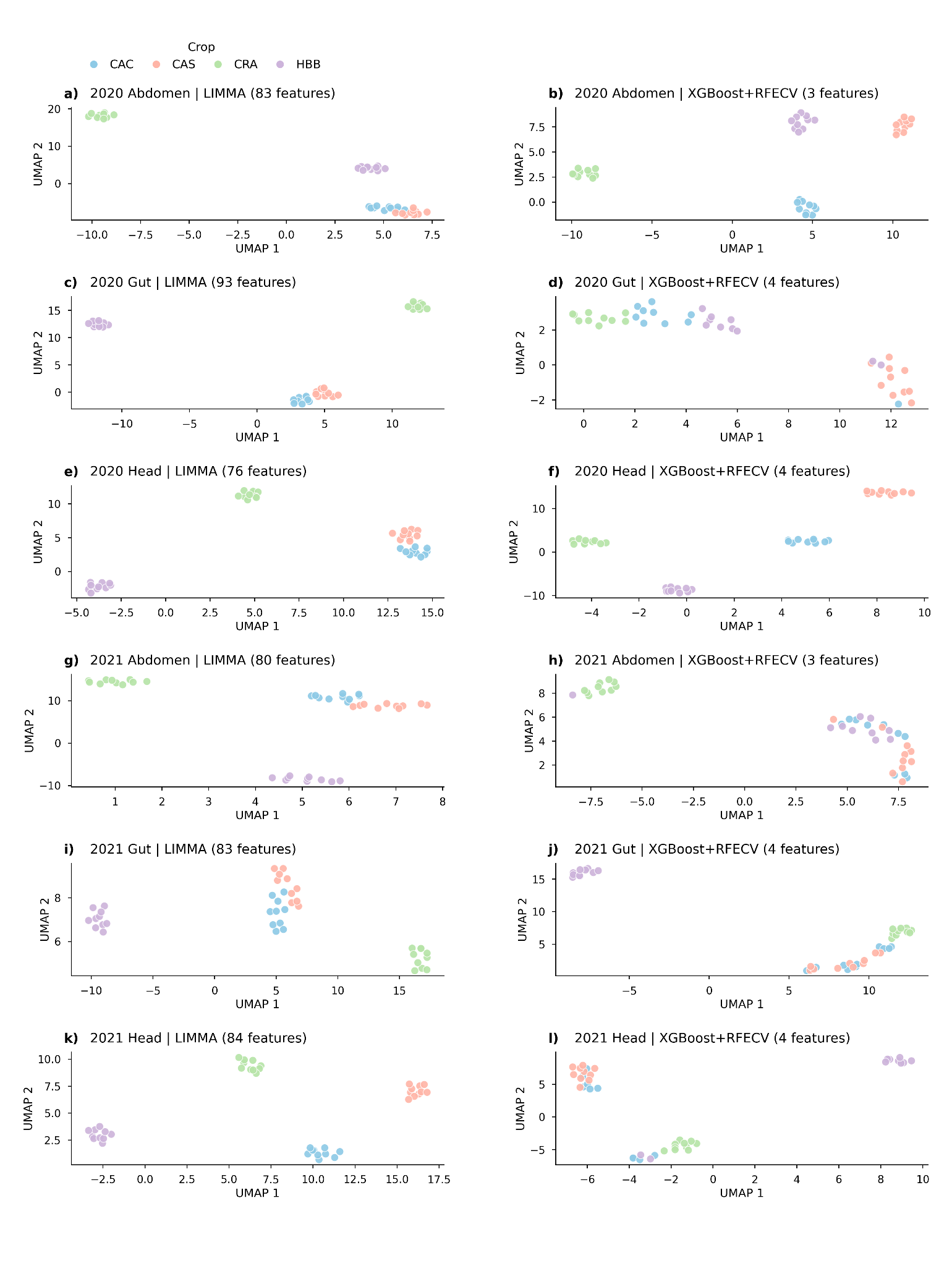

**Supplementary Figure 14. Tissue-year-separated UMAP visualisation of LIMMA-derived descriptive reference features and fixed XGBoost+RFECV protein panels.**

**(a– l)** Each panel shows a tissue-year-specific UMAP embedding coloured by crop group. LIMMA feature sets were constructed separately within each tissue-year context, whereas XGBoost+RFECV feature sets were selected from 2020 only and applied unchanged to 2021. The comparison shows that broad LIMMA references often produce strong descriptive separation, while compact XGBoost+RFECV panels retain crop-associated structure in several tissue-year settings despite using only three to four protein features.

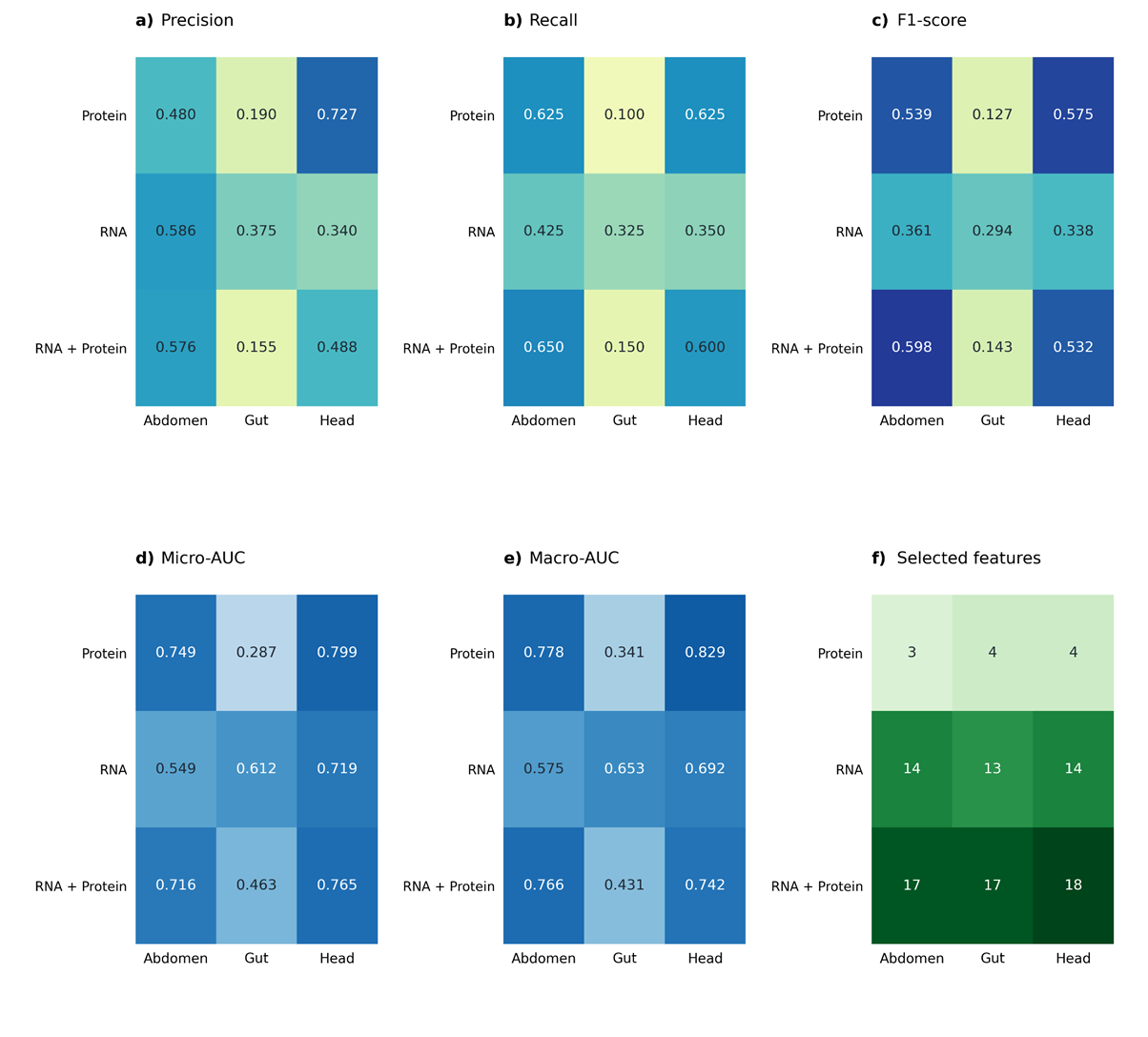

**Supplementary Figure 15. Independent-year prediction performance of XGBoost classifiers trained on RFECV-selected feature panels.**

In every heatmap, rows denote the input feature modality (Protein, RNA, or RNA + Protein) and columns denote the tissue (abdomen, gut, head); cell values are the corresponding metric. **(a)** Precision. **(b)** Recall. **(c)** F1-score. **(d)** Micro-averaged AUROC. **(e)** Macro-averaged AUROC. **(f)** Number of features retained after RFECV. Feature selection and hyperparameter tuning were performed exclusively on the 2020 training cohort, and all metrics were computed on the independent 2021 validation cohort. Protein-only models achieved the strongest classification performance in head tissue (macro-AUROC = 0.83) and more moderate performance in abdomen (macro-AUROC = 0.78), while requiring only 3-4 features. In contrast, protein-based classification in gut was weak and often near or below chance levels (micro-AUROC = 0.29). Gut performance improved when RNA features were included, reaching a moderate macro-AUROC of 0.65. However, combining RNA and protein data did not consistently improve performance beyond that achieved by the best-performing single modality.
